# Insulation between adjacent TADs is controlled by the width of their boundaries through distinct mechanisms

**DOI:** 10.1101/2024.12.25.630322

**Authors:** Andrea Papale, Julie Segueni, Hanae El Maroufi, Daan Noordermeer, David Holcman

**Author notes:** equal contributions (Andrea Papale, Julie Segueni, Daan Noordermeer, David Holcman). Author contributions: D.N. and D.H. designed research; A.P, J.S., H.E.M., D.N. and D.H. analyzed data; A.P., J.S., H.E.M., D.N. and D.H. contributed new analytic tools; A.P., J.S., D.N. and D.H. wrote the paper. Competing Interest Statement: the authors declare no competing interest. Keywords: TADs, boundaries, loop extrusion, CTCF, polymer modeling.

## Abstract

Topologically Associating Domains (TADs) are sub-Megabase regions in vertebrate genomes with enriched intra-domain interactions that restrict enhancer-promoter contacts across their boundaries. However, the mechanisms that separate TADs remain incompletely understood. Most boundaries between TADs contain CTCF binding sites (CBSs), which individually contribute to the blocking of Cohesin-mediated loop extrusion. Using genome-wide classification, we show here that TAD boundary width forms a continuum from narrow to highly extended and correlates with CBS distribution, chromatin features, and gene regulatory elements. To investigate how these boundary widths emerge, we modified the Random Cross-Linker (RCL) polymer model to incorporate specific boundary configurations, enabling us to evaluate the differential impact of boundary composition on TAD insulation. Our analysis identifies three generic boundary categories, each influencing TAD insulation differently, with varying local and distal effects on neighboring domains. Notably, we find that increasing boundary width reduces long-range inter-TAD contacts, as confirmed by Hi-C data. While blocking loop extrusion at boundaries indirectly promotes spurious intermingling of neighboring TADs, extended boundaries counteract this effect, emphasizing their role in maintaining genome structure. In conclusion, TAD boundary width not only enhances the efficiency of loop extrusion blocking but may also modulate enhancer-promoter contacts over long distances across TADs boundaries, providing a mechanism for transcriptional regulation.

**Significance statement:** Topologically Associating Domains (TADs) compartmentalize vertebrate genomes to limit cross-domain enhancer-promoter loops. Our study reveals that boundaries between TADs are diverse genomic entities that range from narrow to highly extended. Using an interdisciplinary approach that combines genome-wide data analysis with biophysical polymer modeling, we find that wider TAD boundaries reduce spurious long-range interactions between neighboring domains. Moreover, we reveal how different boundary components can create this difference in insulating capacity. Our identification and characterization of TAD boundary width and composition suggests they have the potential to regulate the formation of enhancer-promoter loops across TAD boundaries at close and long distance.

---

Hi-C technology has emerged as the gold standard for studying genome-wide 3D chromatin organization [1]. This technology has revealed the existence of Topologically Associating Domains (TADs) in human and mouse cells, which appear as insulated domains within population-averaged chromosome contact matrices [2]. Within TADs, intra-domain contacts are enriched by approximately two-fold compared to contacts with neighboring regions at similar genomic distances [2, 3]. TADs are separated by boundaries, yet the mechanisms underlying their ability to promote intra- over inter-domain interactions remain incompletely understood. A substantial fraction of these boundaries contain binding sites for the CTCF insulator protein, which recognizes a non-symmetric DNA motif [2, 3, 4]. Before the discovery of TADs, the CTCF protein was already known for its ability to block enhancer-promoter communication [5]. Perturbation or reorganization of CTCF binding sites (CBSs) at TAD boundaries can result in the formation of ectopic enhancer-promoter contacts (EP-loops) [6, 7]. Thus, TADs and the CBSs at their boundaries create ”regulatory neighborhoods” that prevent the formation of undesired EP-loops [3, 8]. Other chromatin-associated factors, such as RNA PolII and the MCM complex, are also capable of blocking loop extrusion, but their roles in establishing regulatory neighborhoods remain less well understood [9, 10]. The mechanism by which distal CBSs at TAD boundaries block the formation of EP-loops remains a subject of intense investigation. A key insight into how TADs may function arises from the observed accumulation of the Cohesin protein complex at CBSs [11, 12]. Initially, Cohesin was recognized for its role in entrapping sister chromatids during mitosis. However, this function has been extended to interphase, where the Cohesin complex is proposed to facilitate the organization of intra-chromosomal DNA loops [13].

Biophysical modeling of polymer behavior has become a powerful tool for investigating the mechanisms, dynamics, and heterogeneity of chromatin organization (*e.g.*, [14, 13, 15, 16, 17, 18, 19, 20, 21, 22]). By incorporating chromatin connectivity, the free and confined motion of chromosome fibers, and local fluctuations across multiple time scales, these models can replicate the structure and dynamics of TADs observed in Hi-C and other chromatin conformation technologies [23, 24, 25, 26, 27]. Among these approaches, the Random Cross-Linked (RCL) polymer model [28, 29] has proven particularly effective. In this framework, TADs are structured by cross-linkers, representing the active process of Cohesin-mediated loop extrusion, which is subsequently halted at boundary sites. These models not only reproduce TAD architecture but also provide insights into the dynamic interplay between loop extrusion and boundary blocking, offering a mechanistic understanding of chromatin organization.

Recent *in-vitro* single-molecule imaging studies have confirmed the extrusion capacity of the cohesin complex, supporting the dynamic nature of this process [30, 31]. In contrast, live-cell imaging studies have suggested that CBSs are inefficient blockers of loop extrusion, making them permeable boundaries [32, 33, 34]. Indeed, clustered CBSs can be found at many TAD boundaries, where removal of individual CBSs has a limited impact on the prevention of ectopic EP-loops [35, 36, 37]. The insulation between neighboring TADs can be improved by a sequential blocking of the extruding Cohesin complex at nearby CBSs, thereby creating extended genomic domains where insulation gradually increases [38]. Such sequential blocking leads to a positive feedback loop, which can guarantee long-time persistence of loops [39].

Various modeling studies have explored whether neighboring TADs influence each other’s intra-domain organization [24, 29, 40, 41]. Traditionally, these studies modeled boundaries separating TADs as single, generic contact points. By advancing the Random Cross-Linker (RCL) polymer model [25, 29, 41], we address here how clustering of CBSs improves TAD boundary function [38]. Recently, we showed that the density of intra-TAD cross-linkers modulates intermingling between neighboring domains by inducing a ”fuzzy” behavior of sequences near boundaries [42]. In this study, we investigate how TAD boundary composition—specifically the size of the genomic interval where loop extrusion is blocked and the organization of CBSs within these intervals—affects long-range intra- and inter-TAD organization. To address this, we developed a multi-scale algorithm to classify TAD boundaries from Hi-C data based on the width of their insulation intervals. Using polymer simulations with specific cross-linker distributions at the boundaries, we successfully reproduced the observed diversity in TAD boundary structure, insulating capacity, and CBS organization. By intersecting these boundary categories with CTCF binding data and other chromatin and regulatory features, we uncovered significant differences in their chromatin make-up and how these differences may influence gene regulation. Furthermore, using mean passage time analysis [18, 43], we evaluated the impact of TAD boundary width on global TAD organization. Unexpectedly, we found that wider insulation intervals significantly reduce inter-TAD contacts, even between genomic regions far from loop extrusion block sites. Extended boundaries not only enhance the local separation between neighboring TADs by improving loop extrusion blocking but also reduce spurious intermingling between domains at greater distances. We propose that these direct and indirect effects of boundary width differentially regulate enhancer-promoter (EP)-loop formation across domains. This suggests that TAD boundary composition plays a crucial role in modulating genome organization and transcriptional regulation at multiple scales.

## 1 Results

### 1.1 Strategy to classify TAD boundaries based on the width of the insulation interval

To identify and characterize different categories of TAD boundaries, we used a data-driven computational workflow to determine the width of insulation as outlined in Fig.1. We previously used this approach to identify the optimal parameters to model the genome-wide average of insulation at TAD boundaries in mouse embryonic stem cells (mESCs) [38]. Here, we further develop the optimal calibration of this model and refine its application to characterize different categories of TAD boundaries based on the width of their insulation interval. Briefly, we reanalyzed population-averaged Hi-C data from mESCs (Fig.1-i, [44]), followed by the identification of TAD boundaries using an insulation score (*IS*) approach (Fig.1-ii, [38, 45]). For each genomic bin, the number of interactions in a sliding square of 500 kb is determined, followed by calling of TAD boundaries when local minima in the *IS* reach below a pre-defined threshold. Importantly, this approach identifies boundaries irrespective of the type of neighboring domains. Besides TADs, these may consist of chromatin structures that are not formed by loop extrusion—like the multi-Megabase Hi-C A/B compartments—although in many cases these domains are present as overlapping nested domains with shared boundaries [1, 2, 46].

**Figure 1:**
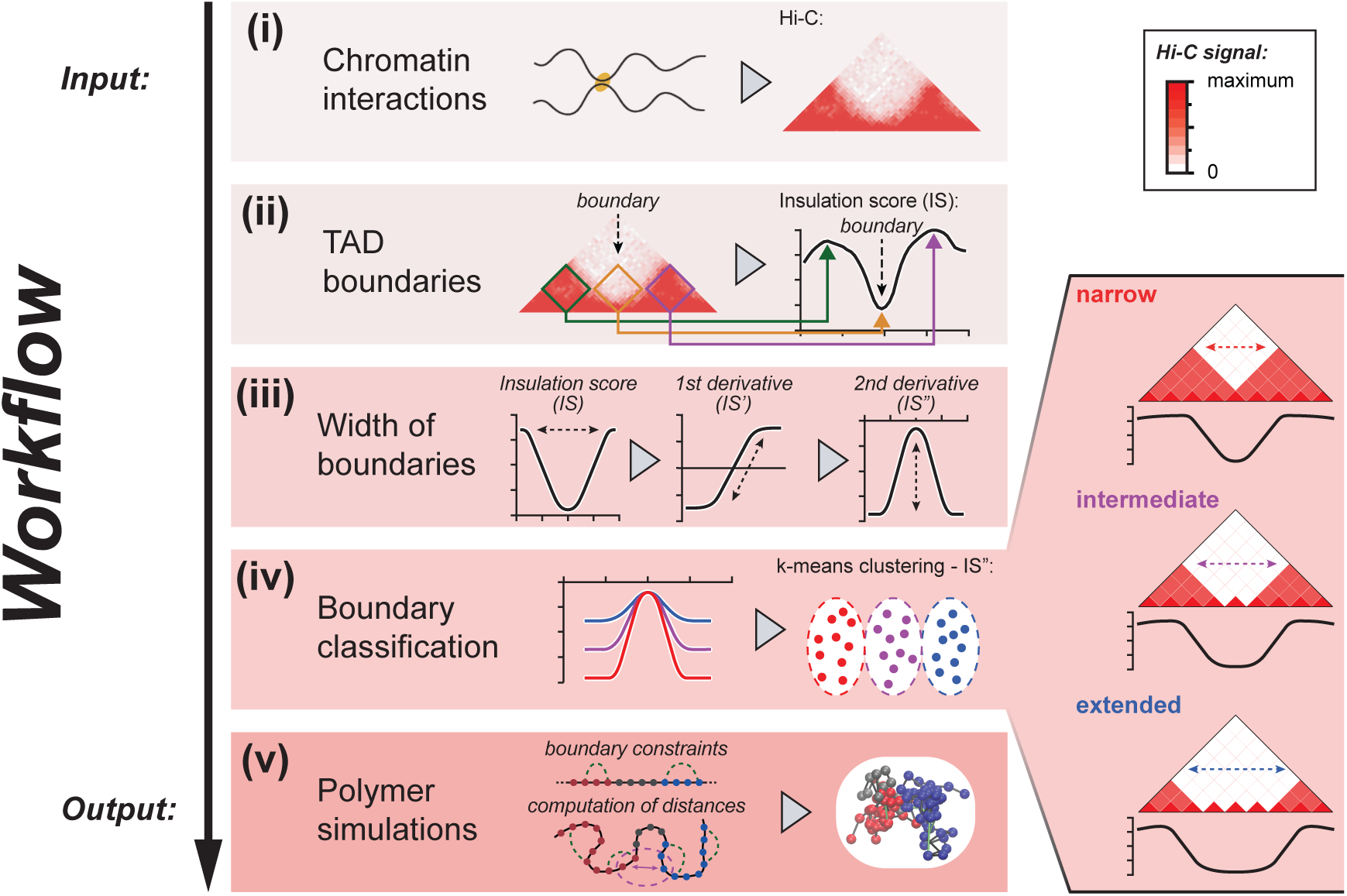
Workflow for TAD boundary classification and quantification. **(i).** Input: genome-wide chromatin contacts are determined from Hi-C data. **(ii).** TAD boundaries are identified using the Insulation Score (*IS*), where cumulative interaction signal is calculated within a moving square along the diagonal in the Hi-C matrix (example: green, yellow and purple squares). TAD boundaries correspond to local minima in the graph (valleys) that reach below a predefined threshold (arrow). Whereas the depth of a valley can be interpreted as the local strength of insulation between TADs, the width of the valley reveals the size of the region that functions as the boundary. **(iii).** Quantification of the width of the boundaries using the second derivative of the *IS* (*IS”*). The depth and narrowness of the *IS”* is a quantitative proxy for valley size. **(iv).** Grouping of TAD boundary size using k-means clustering of the *IS”* allows the identification of three categories: narrow/intermediate/extended TAD boundaries. **(v).** Polymer modeling and simulations with boundary constraints to explore the interface between TADs and the inter-TAD contact dynamics of genomic regions at long distance.

Next, the width of TAD boundaries was ranked by computing the second derivative of the *IS* (*IS”*) with respect to the genomic distance (Fig.1-iii). Using this approach, a narrow TAD boundary is accompanied by a more negative and narrow *IS”* signal within its surroundings, whereas a wider boundary has a less negative and broad signal. To classify TAD boundaries based on the width of their insulation interval, we applied a *k* -means clustering algorithm on the *IS”* with the aim of separating into three generic categories of boundaries (Fig.1-iv). Finally, we developed a procedure to construct the optimal polymer models, whereby the insulation score in the simulations mimicked the experimental data. These polymer models were then used to explore chromatin organization, including about the extent of the boundary regions themselves and about long-range contacts between TADs (Fig.1-v).

### 1.2 Stratification of TAD boundaries in three categories of insulation width

Visualization of 10 kb-resolution Hi-C matrices and their corresponding insulation score (*IS*) from mESCs (obtained after reanalysis of data from [44]) reveals examples of boundaries where the separation between neighboring TADs varies from a sudden drop in the *IS* score (Fig.2A, left) to more intermediate and extended states (Fig.2A, middle and right). Plotting of ChIP-seq data confirms that these boundaries coincide with CBSs and accumulations of Rad21; an essential component of the Cohesin complex that is responsible for loop extrusion. Moreover, in each example, a strong separation between neighboring TADs can be observed at larger distances, confirming they constitute *bona-fide* boundaries. Zooming in on the normalized *IS* for these examples (*i.e.* where the minimum *IS* is set to 0) further illustrates the difference in width of the boundaries (Fig.2B). To systematically classify the 3654 TAD boundaries that we identified in mESCs, we calculated the second derivative *IS”* for all genomic positions, as proxy for the steepness of the *IS* gradient. Subsequent ranking of the *IS”* around each TAD boundary revealed a wide diversity in the width of insulation between neighboring domains (Fig.2C). TAD boundaries ranged, in a continuum, from vary narrow (≈ 10 − 20 kb) to highly extended (≥ 100 kb). The separation between TADs, and in extension the patterns of loop extrusion blocking, are therefore highly diverse and specific to individual boundaries.

**Figure 2:**
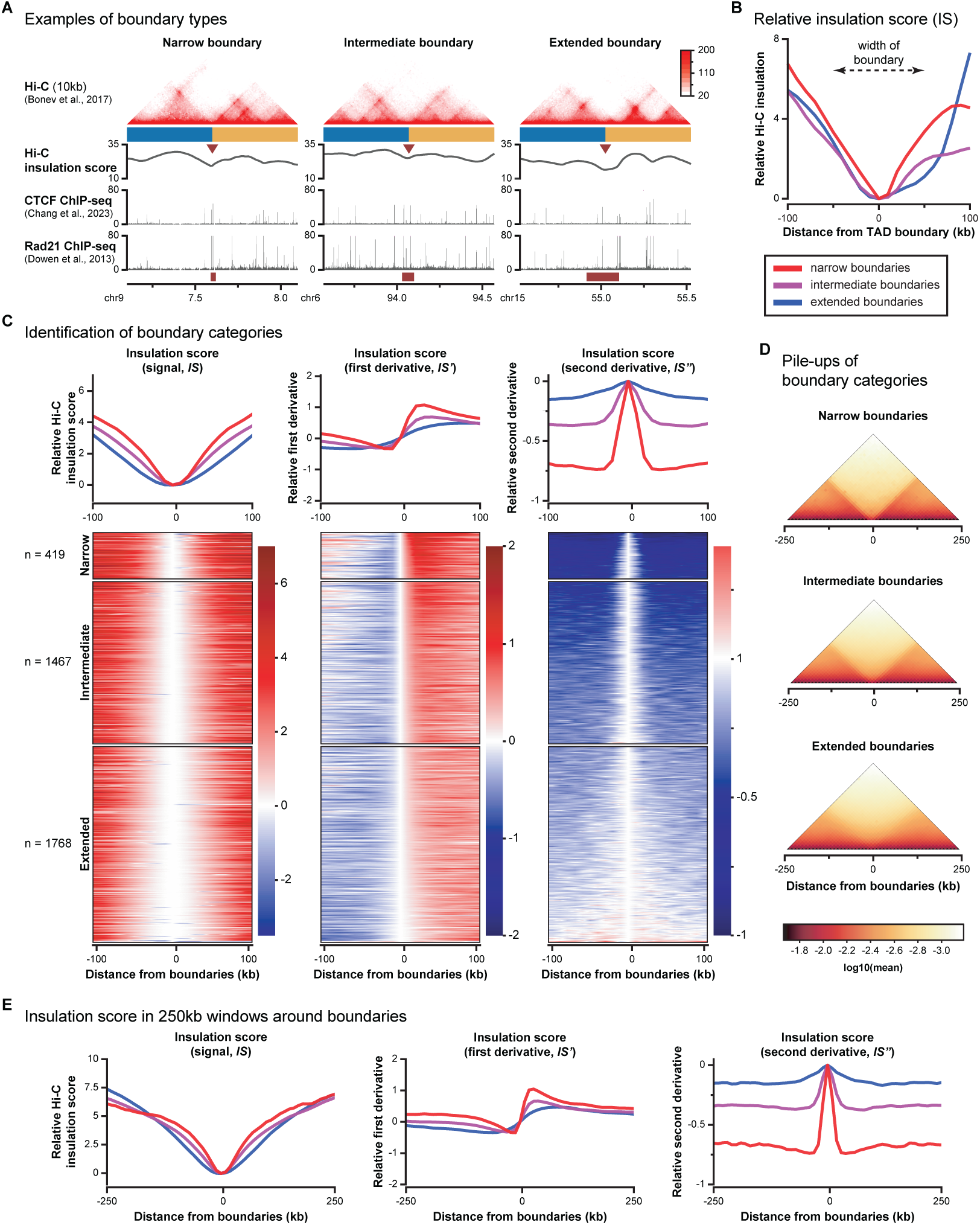
Classification of TAD boundaries. **(A)** Examples of TAD boundaries with different patterns of insulation. Left: a narrow boundary between TADs. Middle: an intermediate boundary. Right: an extended boundary. Hi-C data is depicted above. Purple arrowheads indicate the position of the boundary and the blue and yellow rectangles indicate the up- and downstream domains. Below, tracks for the Hi-C insulation score, CTCF and Rad21 ChIP-seq signal and computationally inferred width of the boundaries (purple boxes; see below) are shown. **(B)** Relative insulation score (*IS*) in a zoomed-in window of +/- 100 kb from the boundaries shown in panel (a). **(C)** *k* -means clustering of TAD boundaries in mouse ESCs, based on the width of *IS* interval. Left: relative *IS* (signal; indicative of the width of the boundary). Middle: normalized first derivative of the *IS* (*IS’*), indicative of the discreteness of the *IS* at the boundary. Right: normalized second derivative of the *IS* (*IS”*), indicative of the steepness of the boundary. Using *k* -means clustering of *IS”*, three categories of insulation score intervals are extracted. Average signal for the three categories in a window of 100 kb up- and downstream of the boundaries is indicated on top and heatmaps for individual boundaries are depicted below. **(D)** Averaged Hi-C matrices for the three categories of TAD boundaries in a window of 250 kb up- and downstream of the boundaries. **(E)** Average *IS* and derivatives for the three categories of TAD boundaries in a zoomed-out window of of +/- 250 kb from the boundaries.

To facilitate downstream analysis, we divided the boundaries into three categories using a *k* -means clustering algorithm (Fig.2C, Fig.S1 and Table S1). Narrow boundaries make up the smallest category (n = 419) and are defined by a sharp transition that coincides with a highly negative and narrow *IS”* peak at the center position (Fig.2C). Intermediate boundaries (n = 1467) are defined by a more extended minimum in the *IS*, coinciding with more intermediate negative *IS”* values, yet a clear transition remains visible in both the *IS’* and *IS”* heatmaps (Fig.2C). Extended boundaries (n = 1768) are defined by a plateau of low *IS* signal around the center position, which coincides with shallow and extended reductions in the *IS”* (Fig.2C).

Whereas the visualization of the normalized *IS* within +/- 100 kb from the boundary may suggest that the three categories are associated with a different degree of separation between the neighboring TADs (Fig.2C, top), a zoom-out to +/- 250 kb reveals that chromatin interactions and the normalized *IS* attain similar values at longer distances, indicating that chromatin organization within and between neighboring TADs on average does not drastically differ between the categories (Fig. 2D,E).

### 1.3 Minimization procedure to construct the optimal polymer representation of the three categories of TAD boundaries

To characterize the three categories of boundaries that we identified, we developed a method to extract polymer models with optimized heterogeneous cross-linker distribution. In these models, a boundary and its two neighboring TADs are represented by two Rouse-polymers of 100 connected monomers with a variable number of randomly positioned cross-linkers (Fig.3A, yellow dashed lines). Essentially, each of these cross-linkers can be considered as the representation of an extruded intra-TAD loop. A boundary is generated by the preferential positioning of cross-linkers within the same polymer (TAD 1: blue monomers 0-100 and TAD 2: red monomers 101-200) over cross-linkers between the polymers [41, 47]. Next, to account for the diversity of TAD boundaries in the Hi-C contact maps (Fig.2), linked to the blocking of loop extrusion within extended genomic intervals, we added further constraints to the distribution of cross-linkers in the polymer (Fig.3B-D). To incorporate the blocking of loop extrusion by CBSs or other chromatin features at the boundary, we fixed cross-linkers at the monomers directly upstream and downstream of the boundary (Fig.3B, green dashed lines). To incorporate the clustering of multiple CBSs within extended boundaries, we included a gap at the boundary that did not interact with the neighboring TADs (Fig.3C, green monomers). To incorporate the variable nature of loop extrusion blocking caused by the inefficient blocking capacity of individual CBSs, we allowed the position of the boundary to move over a defined interval of monomers (Fig.3D, mixed blue/red monomers). Reconstruction of Hi-C matrices from the average configurations of large numbers of simulations for each model (focusing on monomers 50-150) results in little fluctuation, keeping a clear separation between neighboring domains as represented by two discrete ’pyramids’ of strong interaction signal (Fig.3B-D, red signal in simulated Hi-C matrices). Combination of these models and optimization of the parameters for each constraint allow us to increase the accuracy of the separation between the neighboring TADs in the polymer model. In turn, this allows for the reconstruction of different categories of boundaries followed by the determination of monomer behavior at the interface and within the two TADs (see below).

**Figure 3:**
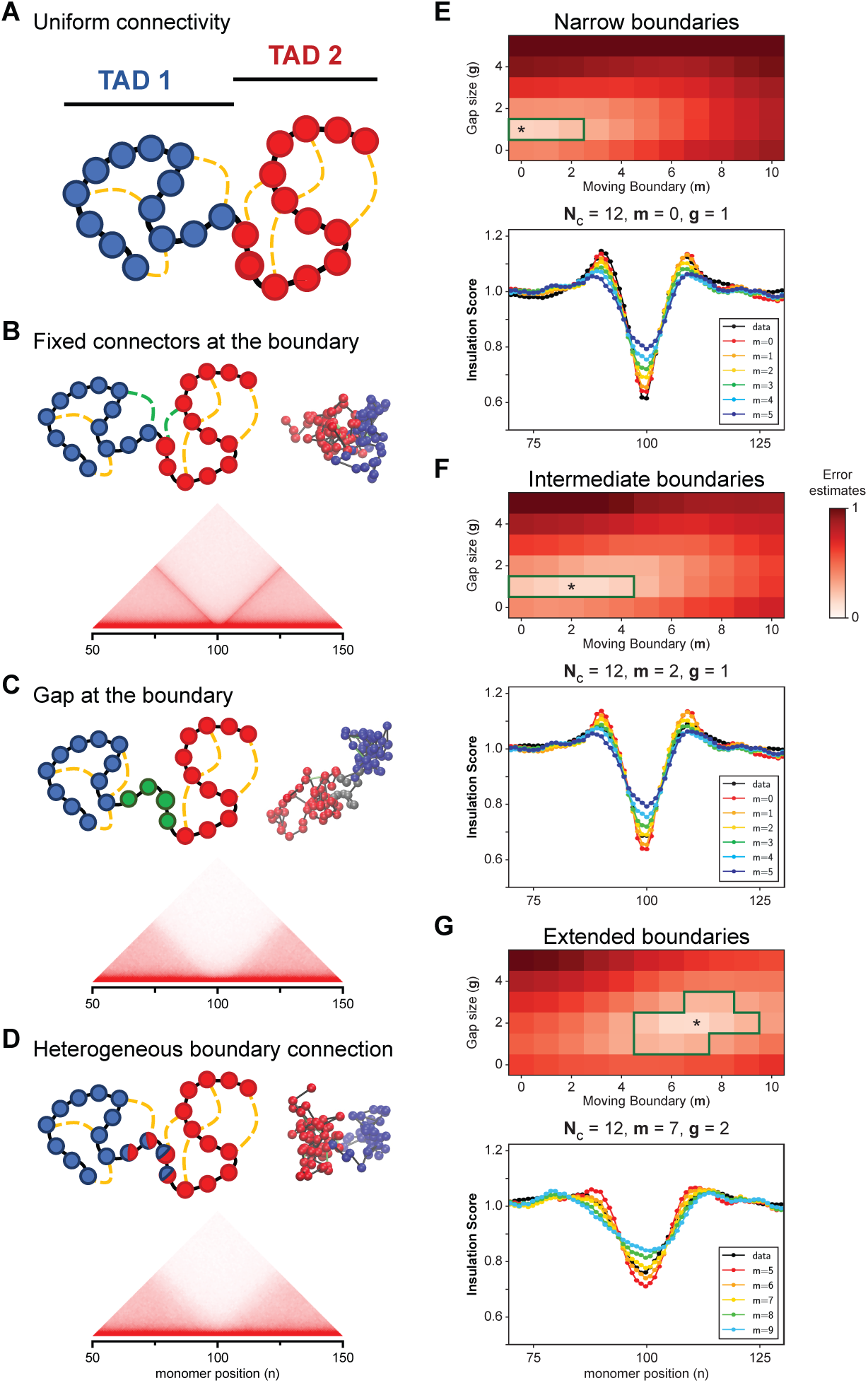
Optimal reconstruction of RCL polymer models to account for the three TAD boundary categories. **(A)** Scheme of polymer representation of two TADs using two concatenated Randomly Cross-Linked (RCL) polymers [41]. Blue monomers belong to TAD 1 and red monomers to TAD 2. Black lines show the backbone connectivity, while yellow dashed lines indicate random connectors with uniform distributions. **(B)** Representation of a polymer with fixed random connectors at both sides of the boundary (connectors in green). **(C)** Representation of a polymer with a gap of *N_gap_* monomers (green monomers; variable *g*) without random connectors separating the two TADs. **(D)** Representations of a polymer with a moving boundary position (monomers with mixed blue/red color; variable *m*). **(E,F,G)** Identification of the parameters that optimally reproduce the *IS* versus monomer location for the narrow, intermediate and extended boundaries.The matrix at the top shows error estimates, with the green box indicating the range of parameters that best reproduce the *IS* obtained from Hi-C data. The * indicates the intersection of parameters where the error estimates are lowest. All models incorporate fixed connectors at the boundary.

A reconstruction of the polymers that best account for the *IS* at the different categories of TAD boundaries first requires the identification of the optimal number *N_mon_* of monomers, the number *N_c_* of cross-linkers, the length of the extended boundary *g* (gap) and the moving *m* of the boundary position. Moreover, the polymers can be optimized with and without the presence of fixed random connectors at the boundary. The variable *g* consists of adding a small polymer chain of a given length without any connectors, whereas the variable *m* accounts for fluctuations among realizations. To identify these parameters, we chose the following criteria: optimal parameters are those that minimize the difference in the *IS* between the simulations based on the polymer model and the experimental Hi-C data, as described by formula:

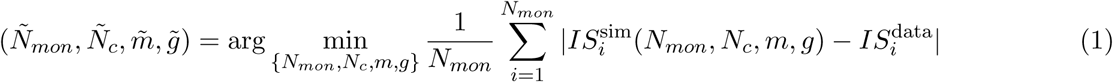

Here, we define the insulation score *IS* as:

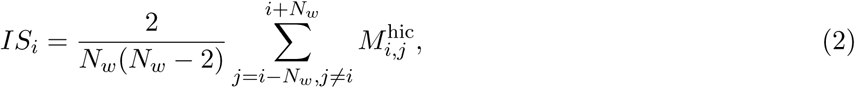

where *M* ^hic^ is the Hi-C matrix (for data) or the Contact matrix (for simulations) while *N_w_* is the window size (see section 3.1.1).

To streamline the procedure for finding the optimal parameters for TAD boudnary composition, we reduced the search in the four dimensional space by first setting the number *N_mon_* of monomers to 100 for each TAD [38]. This results in a polymer that encompasses 200 monomers for the two TADs combined, thereby representing a 2 Mb genomic interval at a 10 kb resolution. Next, we determined the optimal number *N_c_* of cross-linkers, leading to 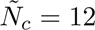, a density that is in a similar range as reported in other studies [24, 32]. Using the reduction to two parameters, we next simultaneously move the position of the boundary within the interval *m* ∈ [0, 10] and varying the size of the gap at the boundary *g* ∈ [0, 5], with the further option of adding fixed cross-linkers at the boundary.

We systematically explored the parameter space to identify the values that best reproduced the experimentally obtained insulation score patterns for the three categories of TADs boundaries. For each set of parameters, we simulated 500 polymer realizations to compute the average contact matrix and the associated *IS*. Using formula (1), we compared numerical simulations to experimental Hi-C data, thus computing the absolute distance between the insulation scores. First, we determined the optimal number *N_c_* of cross-linkers for each boundary category (not shown), using a similar range of values as previously determined [38]. By exploring the space, we consistently identified the minimum for the three boundary categories at *N_c_* = 12: for each category, the optimal solution included having fixed cross-linkers at the boundary and the following parameters: *m* = 0 and *g* = 1 for narrow boundaries, *m* = 2 and *g* = 1 for intermediate boundaries and *m* = 7 and *g* = 2 for extended boundaries (Fig.3E-G and Fig.S2). To conclude, the present minimization procedure reproduces the experimental insulation score and allows to construct polymer models that best represent the three categories of TAD boundaries. These optimal reconstructions can now be used to extract statistical properties of chromatin organization of the TADs that surround the different boundary categories, including monomer behaviour at the boundaries and longer-range intra- and inter-TAD contacts.

### 1.4 Different boundary categories are associated with different chromatin and gene regulatory features

Next, we investigated whether the parameters defining the different boundary categories correlate with distinct distributions of CTCF binding and other chromatin features, and whether these differences are associated with varying impacts on gene regulation. First, using reanalyzed ChIP-seq data [38], we found that CTCF binding is enriched in a large window around all three categories of boundaries (Fig.4A and Fig.S3A; black dashed line versus signal). Yet, for the group of narrow boundaries, CTCF is particularly enriched in a narrow window of around 30 kb surrounding the boundary. For intermediate boundaries, the maximum enrichment is reduced but more gradually decreases over a larger genomic window. CBSs are therefore spread out within a larger region as compared to narrow boundaries. For extended boundaries, the enrichment in CTCF binding is relatively minor and spreads out over an even large domain (Fig.4A and Fig.S3A). Similar patterns are observed for the Rad21 component of the Cohesin complex [48], albeit with a further reduced enrichment around extended boundaries (Fig.4B and Fig.S3B). The distribution and range of CTCF and Rad21 enrichment in the region surrounding the different boundary categories therefore resemble the parameters as extracted using our polymer models.

**Figure 4:**
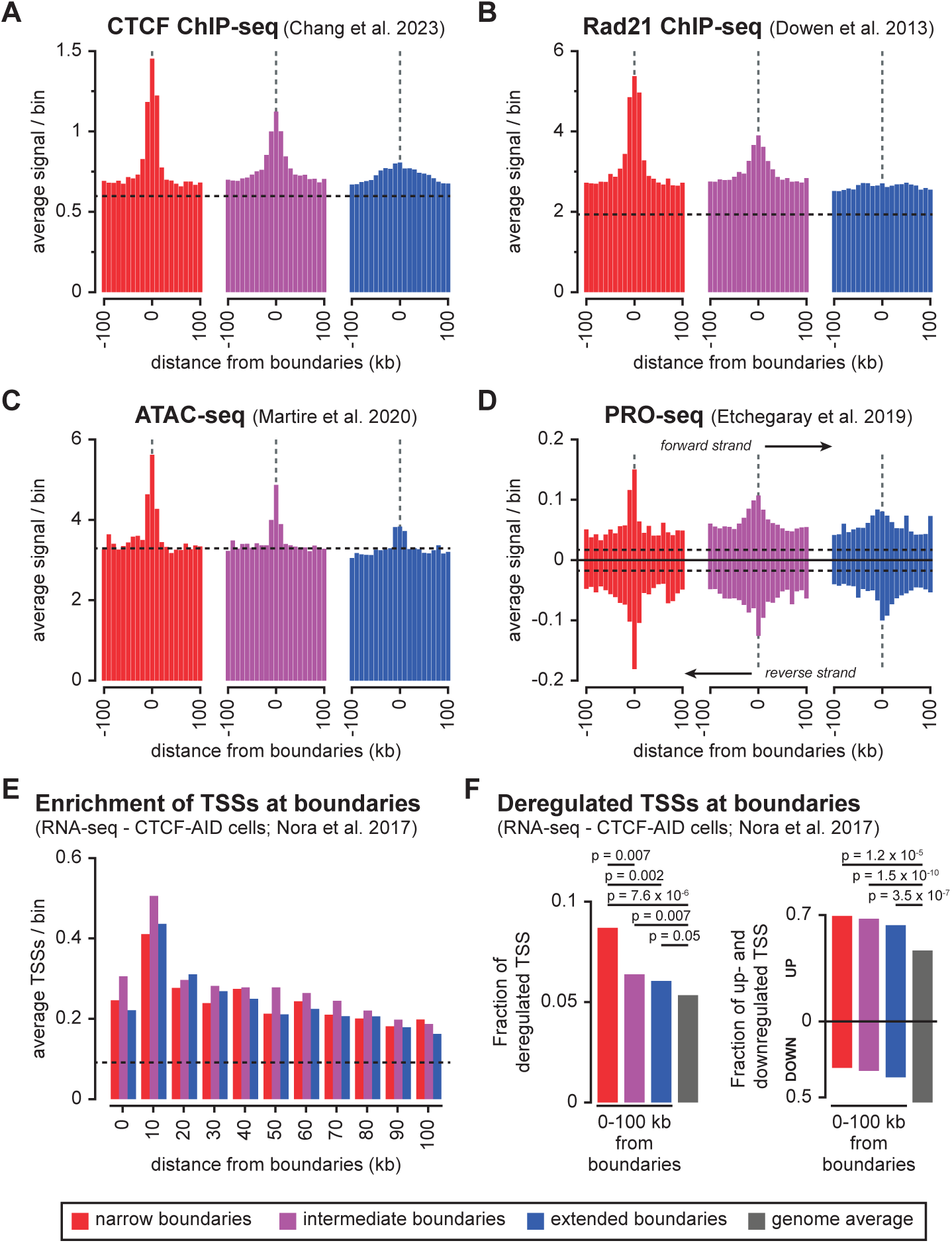
Average enrichment of chromatin and gene regulatory features around the three categories of TAD boundaries. **(A)** Average CTCF ChIP-seq signal. **(B)** Average Rad21 ChIP-seq signal. **(C)** Average accessible chromatin signal (ATAC-seq). **(D)** Average ongoing transcription signal (PRO-seq, with signal separated on the plus and minus strand). **(E)** Average number of TSSs within bins for each TAD boundary category. **(F)** (left) Fraction of deregulated TSS upon CTCF depletion among all TSS mapping within 100 kb from each TAD boundary category (CTCF-AID cells). (right) Fraction of up- and downregulated TSS upon CTCF depletion among all deregulated TSS within the bins (CTCF-AID cells). The origin of each dataset (mouse ESCs) indicated above the panel. Dashed lines represent the average genome-wide enrichment for each feature. Signal in panels **(A-D)** is binned to 10 kb and shown in a window of 100 kb up- and downstream of TAD boundaries. Signal in panel **(E)** is binned to the combined 10 kb up- and downstream of the boundary, and shown in a 100 kb window. Signal in panel **(F)** is binned to the combined 100 kb up- and downstream of the boundary. The gray bar indicates the genome average. Significance of difference between categories was determined using a G-test of independence.

The transcription machinery, including (paused) RNA PolII, has been reported to directly or indirectly influence chromatin insulation as well [10, 49, 50]. We therefore determined the enrichment of accessible chromatin (ATAC-seq), as proxy for active promoters and enhancers, and ongoing transcription (PRO-seq) around the categories of boundaries [51, 52] (Fig.4C,D and Fig.S3C,D). At narrow and intermediate boundaries, both features are comparably enriched as compared to CTCF binding, indicating that they may contribute to the formation of boundaries in these categories as well. In contrast, at extended boundaries, both features are enriched in a narrower window around the boundaries. Combined with the more moderate enrichment of CTCF and Rad21, this suggests that these transcription-associated features contribute more prominently to the formation of this category of TAD boundaries. In line with our biophysical models, the three categories of TAD boundaries are shaped by regions of different size where chromatin features with insulating effects are enriched, and with further specificity for CTCF and other chromatin features.

Having identified the different categories of TAD boundaries and their associated chromatin configurations, we wondered if they could differentially influence gene regulation, for instance through different enhancer blocking efficiencies. To investigate this possibility, we reused existing RNA-seq data from mESCs where rapid CTCF depletion caused a dramatic reduction in insulation between neighboring TADs [53]. In this study, around 5% of genes were called as deregulated after 2 days of CTCF depletion, with roughly equal numbers being up- or down-regulated. Intersection with our boundary categories confirms that the recently reported enrichment of Transcriptional Start Sites (TSSs) close to TAD boundaries is observed for all categories [54] (Fig.4E). Of note, this enrichment is particularly enriched close to the boundary itself (*i.e.* in the bin with the local minimum in the IS, which covers only half the genomic size as compared to the neighboring bins on either side, or the directly neighboring bins at 10 kb distance).

TSSs that are deregulated upon CTCF depletion are significantly enriched within the 100 kb around all three categories of boundaries, which is particularly prominent for the narrow category (Fig.4F, left and Fig.S3E). Moreover, up-regulated TSSs are particularly enriched in all categories (Fig.4F, right). CTCF binding at all categories of boundaries thus preferentially represses TSS activity, which suggests an enriched involvement in the blocking of EP-loops. Interestingly, when investigating these results at higher resolution, a significant enrichment of downregulated TSSs can be observed in the bins that directly overlap with narrow and extended boundaries (Fig.S3F). Whereas this group only includes a small number of TSSs (6 for narrow boundaries and 12 for extended boundaries), this may suggest that CTCF binding for this subset of TSSs has an activating function, for instance by structuring EP-loop formation [55].

### 1.5 CTCF binding is enriched throughout the boundaries but with category-specific differences

Next, we wanted to obtain a more detailed view of CTCF binding within the different TAD boundary categories. To address this question, we first developed a strategy to infer the width of individual boundaries, using the relative *IS”* as input (Fig.5A). Despite fluctuations in the *IS”* value around individual boundaries, each category returned a distinct distribution for the boundary width (Fig.5B and Table S1).

**Figure 5:**
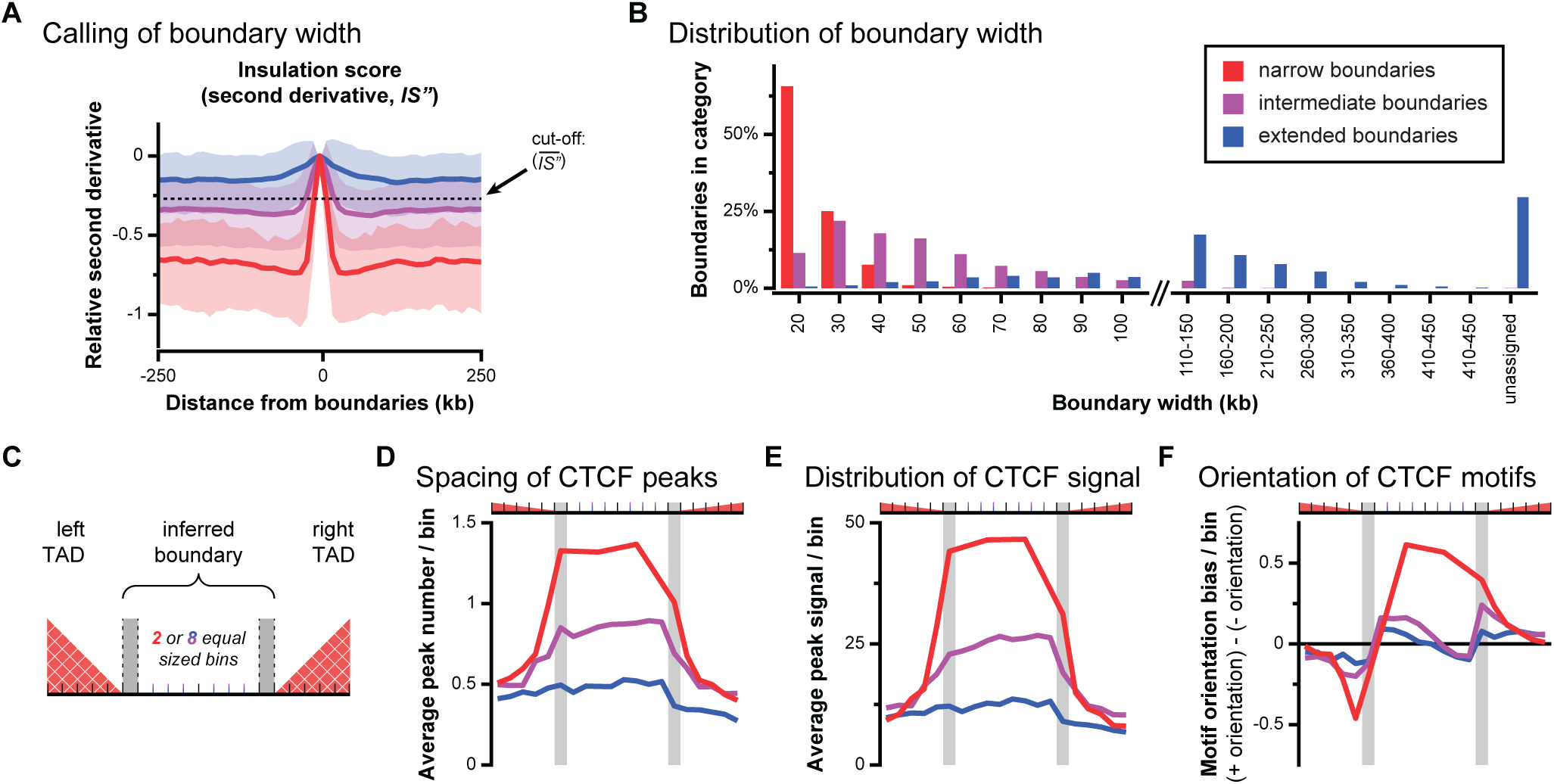
Enrichment of CTCF binding features within inferred boundaries. **(A)** Outline of the strategy to infer the width of individual boundaries using the relative *IS”*. Individual boundaries extend from t<u>he ce</u>nter bin up to the first bin on either side that has a value below a cut-off that is defined as the average relative *IS”* value (*IS*”) within the 250 kb up- and downstream of all boundaries that were included in this study (dashed line). Coloured lines represent the average *IS”* signal (10 kb bins) for each category of boundaries, with shaded areas indicating the 10%-90% interval of the signal. **(B)** Histogram with the distribution of inferred width for individual boundaries in the three previously identified categories. **(C)**Visualization strategy for the spacing of CTCF features within and around extended boundaries. For all boundaries, signal in the two bins at the extremities (grey bins) and 5 first bins in the left and right TAD are determined. For boundaries that span 3 or more bins, the internal bins are proportionally divided over 2 (narrow boundaries) or 8 bins (intermediate and extended boundaries). For all resulting bins, the signal is normalized to the number of boundaries for each category, or to the boundaries that span 3 or more bins (for signal within the boundary). **(D)** Average number of CTCF peaks per bin within and around the inferred boundaries. **(E)** Average CTCF peak signal per bin within and around the inferred boundaries. **(F)** Average CTCF motif orientation bias per bin within and around the inferred boundaries. Bias is determined by substracting the average number of motifs in a positive orientation from the number of motifs in a negative orientation.

For narrow boundaries, over 65% of boundaries span 20 kb, with nearly no boundaries that cover more than 50 kb. The large majority of boundaries in the intermediate category range from 20 kb to 100 kb (median: 40 kb, mean: 49 kb). For extended boundaries, we could assign a width for 1245 out of 1768 boundaries (see Material and Methods section), with a broad distribution of inferred widths (median: 130 kb; mean: 150 kb). Overall, the distributions of inferred lengths align well with the range of parameters in our biophysical models for the different boundary categories (combined *m* and *g* parameters, 3E-G). To characterize different features of CTCF binding within the inferred TAD boundaries, we intersected their individual span with an available list of CTCF peak features in mESCs [38]. To obtain an average description of CTCF feature spacing around the individual boundaries, we used a visualization strategy as described in (Fig.5C). Visualization of CTCF peak spacing and the distribution of CTCF peak signal reveals an elevated signal within the boundaries for the narrow and extended categories as compared to the immediate surroundings (Fig.5D,E). CBSs can thus be enriched throughout the entire genomic interval covered by the boundaries (as compared to only at the extremities), which for intermediate boundaries can range up to considerable distances. A further analysis of CTCF peak signal reveals that narrow and intermediate boundaries do not only differ in peak density (Fig.5D), but that peak signal within intermediate boundaries is globally reduced as well (Fig.S4). In contrast, for the extended category of boundaries a more diffuse pattern of CTCF distribution is observed, supporting the notion that other chromatin features may play a more prominent role. Next, we also investigated if boundaries were associated with a different bias in CTCF motif-orientation (Fig.5F). As expected, CTCF motifs at the extremities or immediately outside of the boundaries were preferentially orientated away from the boundary (see *e.g.* [56, 57]). Interestingly, within the boundaries in the intermediate and extended categories of boundaries, we observed a bias for a convergent orientation of CTCF motifs (*i.e.* motifs that are oriented away from the nearest boundary; Fig.5F). A subset of these boundaries may thus be organized into domains that share similarities with TADs, but whose insulation is insufficient to appear as separated boundaries (for example, see the extended boundary in Fig.1A). Further supporting our biophysical models, this analysis confirms that insulation can spread out all over the regions that are covered by the boundaries.

Having formally confirmed the enrichment of CTCF binding throughout TAD boundaries, we wondered if boundary width could become reorganized during cellular differentiation. We thus reanalyzed Hi-C data from Neural Progenitor Cells (mNPCs) and Cortical Neurons (mCNs), originating from the same study as the mESC data we used in this study [44]. Visualization of the normalized *IS* and *IS”*, similar to (Fig.2C), revealed highly similar distributions and averages for both measures (Fig.S5A). TAD boundary width therefore appears not drastically remodeled during cellular differentiation. Visual inspection of the *IS* nonetheless reveals rare examples of loci where TAD boundary width is reorganized, which coincide with locus-wide changes in CTCF binding (Fig.S5B). Notably, this includes the *Zfp42* pluripotency gene—known as *REX-1* in human cells [58]—where an extended TAD boundary that is marked by multiple CBSs in mESCs is erased in CNs (Fig.S5B, right).

### 1.6 TAD intermingling defined by overlapping radius of gyration

Currently, there are no generic approaches to characterize intermingling between neighboring TADs. Using the outcomes from our biophysical modeling, we can use the gyration radius to determine the degree of intermingling at the interface between TADs. Essentially, the gyration radius represents the average distance from the center of mass of the TAD, as determined by averaging all realizations of the polymer model (Fig.6A). By determining the overlap between the spheres created by the gyration radii from the two TADs, we can determine the intermingled monomers at the interface (Fig.6A). Indeed, each TAD can be delimited by a sphere centered at the centers of mass (CM) using the formula:

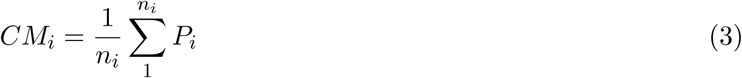

**Figure 6:**
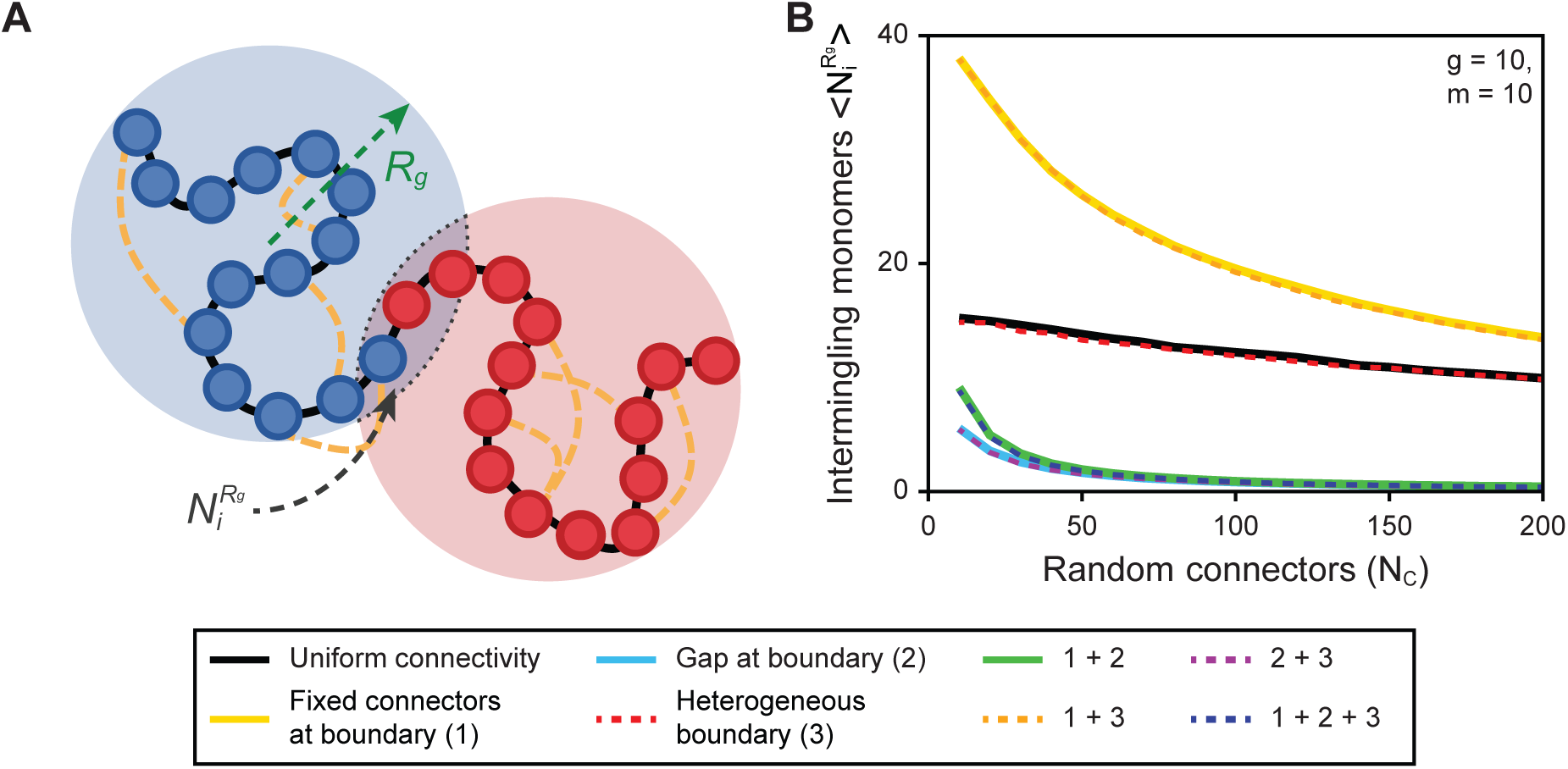
Effect of TAD boundary components on the gyration radius and TAD interface intermingling. **(A)** Two polymers connected by boundaries incorporating additional constraints. Intermingling at the interface (blue-red shading) is defined by the intersection of the two spheres at the center of mass with a radius equal to the radius of gyration. **(B)** Intermingling at the interface measured by the number of monomers that occupy the overlap between the spheres. Colored lines represent the combinations of added boundary components, relative to the number of random cross-linkers *N_c_*.

The radius is the gyration radius:

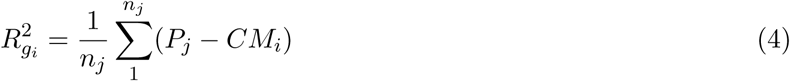

In these formulas, *n_i_* is the index of monomers in each TAD (i=1,2). TADs are then generated by adding randomly distributed connectors.

To determine if the TAD boundaries can have an impact on the overlap of the gyration radius, we use the three additional components that were defined in (Fig.3B-D): (1) adding fixed connectors at the boundary, (2) including a gap at the boundary by adding a small Rouse polymer and (3) creating a heterogeneous boundary by varying the last monomer that belongs to a given TAD (Fig.6B). To obtain more prominent differences, values *m* = 10 and *g* = 10 were used for simulations. Models based on uniform connectivity, *i.e.* without additional boundary components, display a degree of intermingling that is relatively refractory to the number of random connectors *N_c_* (black line). Unexpectedly, the addition of fixed connectors or the case of fixed connectors with a heterogeneous boundary result in a strong increase in the number of intermingling monomers at the interface, particularly at low *N_c_* values (Fig.6B; yellow and orange lines). The incorporation of the parameter that represents loop extrusion blocking thus actively promotes contacts in the region surrounding the boundary. This increase extends over the full range of *N_c_* values, which represents the density of extruded loops, indicating that this intermingling is not directly dependent on the loop extrusion process.

Interestingly, an opposite statistical jump can be observed for all models that incorporate a gap at the boundary (through the addition of a small polymer between the two TADs. This gap represents the presence of multiple CBSs within extended boundaries in our model (Fig.6B; light blue, green, purple and dark blue lines). Here, the number of intermingling monomers is strongly reduced and reaches nearly 0 at increased *N_c_* values. Although the presence of fixed connectors counters this effect at lower *N_c_* values (green and dark blue lines), intermingling remains considerably reduced as compared to a uniform connectivity (black line). Besides their impact on loop extrusion blocking, arrays of CBSs at TAD boundaries—as represented by the gap—thus reduce the intermingling at the interface between the TADs. Finally, the addition of heterogeneity in our models, by incorporating a variable boundary position, did not noticeably impact the degree of intermingling (Fig.6B; red, orange, purple and dark blue lines) despite its important contribution to the reproduction of the local *IS* (Fig.3E-G and Fig.S2).

### 1.7 Impact of boundary components on inter-TAD Mean First Encounter Time

The radius of gyration analysis provides an estimate of the average number of monomers at the immediate interface between two TADs at steady-state (Fig. 6). However, this approach does not account for the dynamic and transient behavior of the chromatin fiber, including the possibility that loci at further away from the boundary may still interact at lower frequencies. To better understand how boundary components influence the dynamic insulation between neighboring TADs, we performed a Mean First Encounter Time (MFET) analysis. In this approach, the mean time is computed that it takes for two monomers within the same or neighboring TAD to come within short distance (as defined by a ball of radius *ɛ*) (Fig.7A). This parameter is particularly relevant for regulatory elements in neighboring TADs, as it can reveal the potential for enhancers and promoters to engage in spurious inter-TAD contacts.

**Figure 7:**
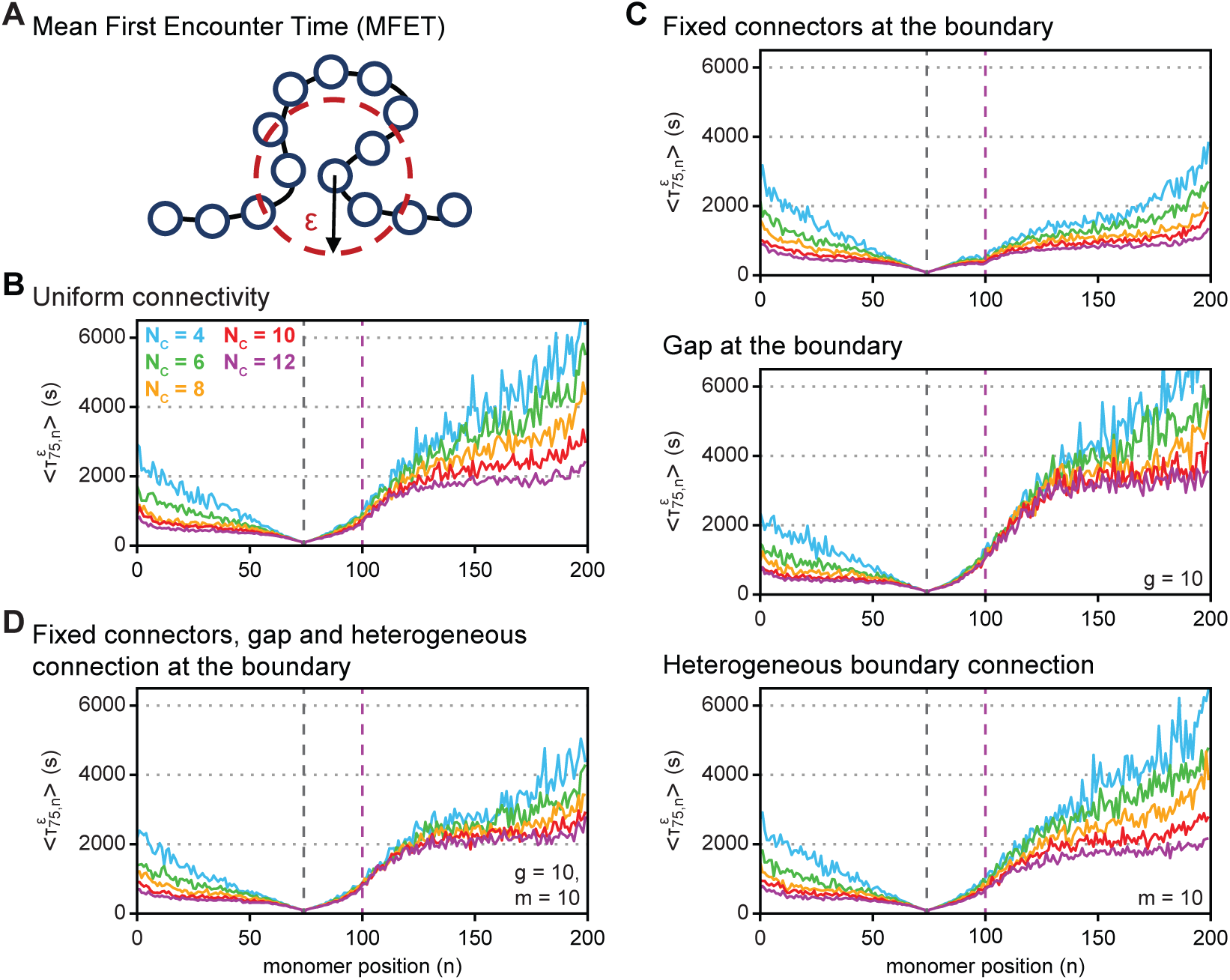
Mean first encounter time (MFET). **(A)** Scheme for the mean first encounter time (MFET), defined as the first interaction time when the distance between two loci becomes smaller than a length *ɛ* between two monomers. **(B)** MFET 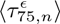 for the uniform connectivity (*i.e.* two TADs defined only by random connectors with uniform distributions) with different numbers of random connectors *N_c_* = 4*, ..,* 12 obtained by averaging over 1000 polymer realizations. The black dotted line indicates the position of monomer 75 and the purple dotted line the boundary between TAD 1 (left) and TAD 2 (right). **(C)** MFET for connectivities with a fixed connector at the boundary (top), with a gap at the boundary (middle) and a heterogeneous boundary connection (bottom). **(D)** MFET for combined connectivities (fixed connector, gap and heterogeneous boundary).

The MFET is determined for all pairs of loci in the two TADs [59, 16]. While the MFET for two monomers in a Rouse chain [60, 59] depends on the polymer length and the diffusion coefficient only, in the RCL-polymer it further depends on the number of cross-linkers [28]. In general, the MFET depends on the local chromatin connectivity [41] and can only be extracted from numerical simulations. Until now, the MFET for loci positioned in neighboring TADs has not been reported relative to boundaries with different components. We thus explored for a given monomer *m*, how the MFET evolves with the genomic distance in the immediate neighborhood of the monomer *m*. For this purpose, we first compute the first encounter time 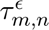 for a single realization between monomers *m* and *n* and then compute the mean (MFET) from large numbers of realizations. By definition, the first encounter time 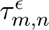 is the first time when two monomers *m* and *n* enter within a ball of radius *ɛ* = *b/*3 (Fig.7A). The ensemble average 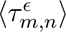 is determined over 1000 polymer realizations. We focus our analysis on monomer 75, which is the center of the 51-100 interval that is the focus of our attention in TAD 1 (Fig.S6A).

Using models with uniform connectivity, *i.e.* 2 TADs without additional boundary components, the MFET reaches a plateau after a few monomers that depends on the level of connectivity (Fig.7B for number of cross-linkers *N_c_* = 4*, ..,* 12 and Fig.S7B for more exaggerated models with a focus on monomer 25 and a larger range *N_c_* = 10*, ..,* 90). Notably, two different plateaus can be observed: a lower value MFET with monomers in TAD 1 (left) and a high value MFET with monomers in TAD 2 (right) (Fig.7B). The transition across the boundary spans around 20 monomers, which represents 10% of the polymer length. Increasing the number of cross-linkers decreases the absolute difference between the MFET in the left and right TADs. However, the number of contacts between the two TADs remains 2-3 fold lower as compared to the steady state within TAD 1.

Next, we determined the impact of adding further constraints to the boundaries. When fixed connectors are added to the boundary, thereby simulating the blocking of loop extrusion, we observed an unexpectedly drastic reduction of the MFET in TAD 2 (Fig.7C-top; monomers 101-200 and Fig.S6 and S7). Similar to our analysis of boundary intermingling (Fig.6B), this effect is independent of the level of connectivity (*i.e.* number of cross-linkers). The addition of simulated loop extrusion blocking to the model thus promotes contacts at larger distance from the boundary as well. Conversely, upon the addition of a gap between the two TADs, the transition of the MFET between the two TADs is considerably increased and remains high when additional connectors are added (Fig.7C-middle and Fig.S6 and S7). The presence of arrays of CBSs to create a more extended boundary thus strongly reduces spurious long-range inter-TAD contacts, similar to the boundary intermingling as well. Importantly, this difference among models is consistent for the same number of cross-linkers, indicating that it is a direct effect of adding a gap at the boundary, rather than differences in polymer organization due to the simulated formation of intra-TAD loops. To further explore the consequences of heterogeneous separation, we introduced a moving boundary as well (Fig.7C-bottom and Fig.S6 and S7). Whereas this parameter had an important impact on the local *IS* pattern (Fig.3E-G), we found that the MFET is equivalent to the model of uniform connectivity within the 2 TADs (Fig.7B).

The combination of the boundary characteristics reveals a differential contribution: adding fixed connectors or a gap dominates over the moving boundary (Fig.7D and Fig.S6 and S7). In contrast, the addition of both fixed connectors and a gap results in a more intermediate inter-TAD MFET value, incorporating the effect of both parameters (Fig.7D and Fig.S6 and S7). Interestingly, in this latter model the MFET remains mostly stable over the *N_c_* range, attaining similar values as compared to the 2 TADs with a uniform cross-linker distribution when the *N_c_* = 12 value is used that we extracted from our simulations in (Fig.4). Compared to models with fixed connectors alone, representative of narrow boundaries, we find that the addition of parameters that mimic wider boundaries lead to an increase in the inter-TAD MFET values by around three-fold.

In summary, besides their involvement in the local blocking of loop extrusion—thereby improving the reconstruction of the local pattern of insulation score *IS* (Fig.4)—we find that the different boundary components have an unexpected influence on more spurious long-range inter-TAD contact probabilities as well (Fig.6 and 7).

### 1.8 Determining TAD boundary extent using the Mean First Encounter Time

Analogous to the radius of gyration (Fig.6), the MFET can also be used to determine the extended nature of boundaries and the behavior of the interface between TADs, while incorporating the dynamic nature of chromatin organization as well. To characterize intra-TAD contacts and their deviation at the boundary, we use the following parameters:

- **Mean encounter time between two locus** *m* **and** 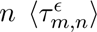 which is obtained by averaging over polymer realizations for the same random connector configuration.
- **Mean encounter time across a TAD** 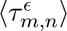, which is computed by averaging 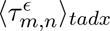 over the monomer *n* belonging to TAD *x* = 1, 2.
- **Mean encounter over a TAD** 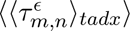, which is computed by averaging 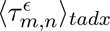 over *m*, for all monomers of the polymer.
- **Mean encounter gap** 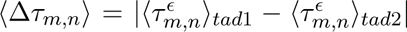 which represents the difference of the MFET between the two TADs, thus measuring the asymmetry of encounter from one monomer in one TAD to a monomer in the other TAD.

To quantify how a centrally located locus (here the monomer at position 50) interacts with its own (TAD 1) and the neighboring TAD (TAD 2), we computed the mean time 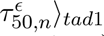 and 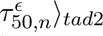. The locus shows a constant level of contacts within its own TAD (yellow and orange bars) but has a strong increase (factor 10) when passing the boundary (Fig.8A; between n=100 and 101). However, when considering a locus just left of the boundary (n=100), interactions were more evenly spread out over the both TADs (Fig.8B). Finally, we computed the map for the encounter time of locus *n* = 150, showing the mirrored situation of case *n* = 50 (Fig.8C).

**Figure 8:**
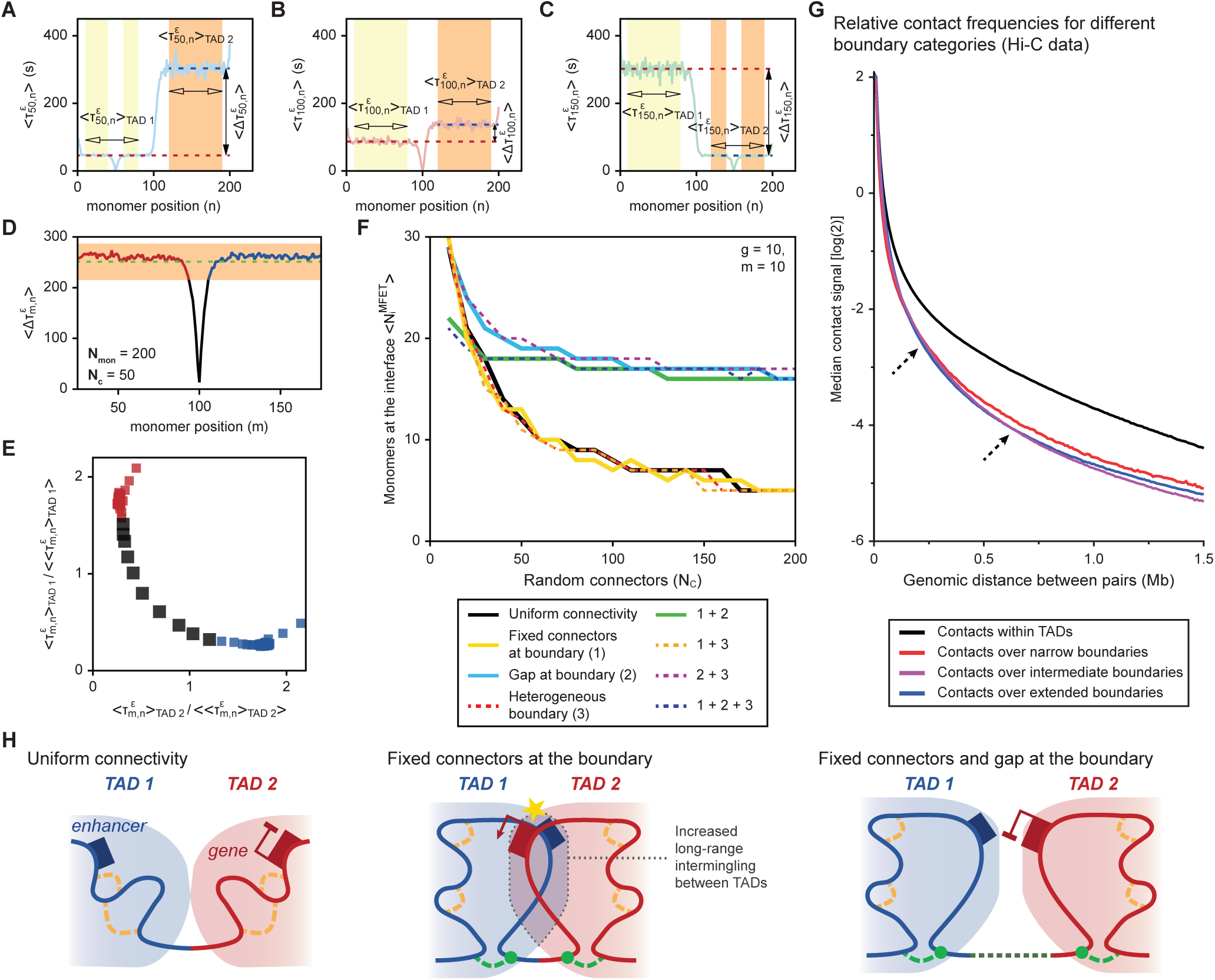
Boundary interface defined by Average Mean Encounter. **(A-C)** Two TADs [1–100] and [101–200] are interacting with different type of connectors. Taking n=50, 100 and 150 as reference monomer. The three panels illustrate how mean encounter time 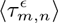 varies with the index position *m*. The mean encounter times 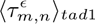 (orange) and 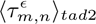 (yellow band) change drastically when passing the boundary, showing how the boundary affects the encounter time between the two TADs. **(D)** Determination of the boundary interface using the average mean encounter. Monomers belonging to a given TAD (red or blue) show a nearly constant average mean encounter. A monomer belongs to the interface when this value falls outside a threshold defined by the standard deviation standard (orange) from the average (dashed green line). **(E)** Visualization of the classification depending on 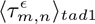 and 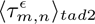 **(F)** Boundary interface defined by the mean number of monomers 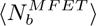 (black) versus number of connectors *N_c_* for the different classes of connectivity. **(G)** Genome-wide Hi-C contacts for pairs of loci at varying genomic distances (10 kb resolution). Pairs of loci are classified for being within a TAD or being separated by a TAD boundary from one of the three categories. Arrows indicate genomic distances where boundary categories intersect. **(H)** Cartoon summarizing the impact of TAD boundary configuration on long-range TAD intermingling. The chromatin backbone in the TADs is indicated with the blue and red solid lines. Random connectors, representing extruding DNA loops, are indicated with yellow dashed lines. Fixed connectors, representing DNA loops that are blocked at the boundary, are indicated by green dashed lines. The potential for ectopic inter-TAD EP-loop formation is illustrated by the presence of an enhancer in TAD 1 and a promoter in TAD 2. The yellow star indicates the formation of a productive loop.

To quantify the insulation for a given monomer *n*, we use the difference of encounter time 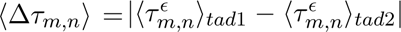, which quantifies an abstract energy barrier for a locus in one TAD to interact with any locus in the other TADs. As an illustration, we plotted this energy barrier for the boundary locus (n=100) when there are 50 connectors in each TAD (Fig.8D). This analysis can next be used to define the extended nature of the TAD boundary: the standard deviation of the mean encounter time is computed (Fig.8D, orange shading) and centered around the mean (dashed green line). Loci outside the orange band are considered to be part of the interface at the boundary (black boxes in Fig.8E). Using this approach, we then compute the number of loci at the interface after adding different boundary components and varying the number of cross-linkers *N_c_* (Fig.8F). Using this approach, three groups emerge:

1. A first group with a gap at the boundary and with or without heterogeneity (Fig.8F; light blue and purple lines). Here the boundary interface is large at low *N_c_* numbers and reaches a plateau of boundary size 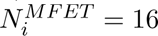 at higher *N_c_* numbers.
2. A second group that combines fixed connectors with a gap at the boundary (Fig.8F; green and dark blue lines). Here the size of the boundary interface is less extended at low *N_c_* numbers 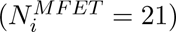 but remains mostly stable with increased *N_c_* numbers.
3. A third group that contains all models without a gap at the boundary (Fig.8F; black, yellow, red and orange lines). Here the boundary interface is large at low *N_c_* numbers but reaches a low plateau of 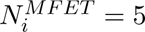 at higher *N_c_* numbers.

Despite similarly grouping the different boundary components together, the gyration radius and MFET analyses return different estimates for the boundary size, particularly at higher *N_c_* numbers (Fig.6 and Fig.8). Yet, at lower *N_c_* numbers, which optimally reproduce local insulation around the boundary (Fig.3E-G), these differences are less pronounced. To conclude, instead of defining TAD boundaries as fixed genomic intervals that create insulation, our First Encounter Time analysis reveals that the different boundary components influence the boundary interface in a more dynamic and variable manner.

To confirm the outcomes of our polymer simulations, we determined the distance-dependent Hi-C contact signal for the different boundary categories in mESCs (Fig.2C and Fig.8G). Compared to intra-TAD pairs of loci (black line), the presence of any type of TAD boundary reduces contact signal by about two-fold over a wide range of distances (one log(2) difference; see also [3]). Yet, differences for the boundary categories can be observed as well. Narrow boundaries create an elevated separation between loci in the neighboring TADs at short range (≤ 100 kb), as may be expected from to more discrete nature of the boundary. Conversely, at intermediate distances (≥ 200 kb), inter-TAD contacts are increased between loci that are separated by narrow boundaries as compared to loci separated by intermediate and extended boundaries. At distances of (≥ 500 kb, this difference reaches up to 1.2 fold between narrow and intermediate boundaries, constituting a non-negligible difference at the genome-wide scale (Fig.8G). Extended boundaries display a similar difference as compared to narrow boundaries, although less pronounced at distances (≥ 500 kb. TAD boundary width thus directly reduces inter-TAD contact levels, whereby intermediate boundaries particularly improve separation over long distance (Fig.8G,H and discussion).

## 2 Discussion

TADs and their boundaries have emerged as major features of mammalian genome organization and the regulation of EP-loops [2, 3]. Previous studies have reported that TAD boundaries can extend over wider ’zones of transition’ that incorporate multiple instances of CTCF binding and that this can influence inter-TAD EP-loops [3, 35, 36, 37, 38, 61]. Yet, a genome-wide analysis to determine the diversity of TAD boundary organization and how this influences local and long-range inter-TAD contacts had not been reported.

In this study, we show that the ≈ 3500 TAD boundaries in mouse embryonic stem cells form a continuum that ranges from narrow (20 kb) to highly extended (≥ 100 kb) zones dedicated to the blocking of loop extrusion (Fig.2). Using optimized cross-linker based polymer simulations, we show that incorporation of parameters that mimic multiple instances of variable loop extrusion blocking allow the reproduction of TAD boundaries with different widths. Intersection of these categories of TAD boundaries with other chromatin and transcriptional features, in combination with a new approach to infer the span of individual boundaries from the Hi-C matrix, confirms that narrow boundaries coincide with a highly focal enrichment of CTCF binding, Cohesin accumulation and sites of ongoing transcription. Conversely, more extended boundaries coincide with wider and less pronounced patterns of enrichment for these chromatin features (Fig.4 and Fig.5). Moreover, we confirm the enhancer blocking effect of all boundary categories on nearby genes, which is particularly prominent for narrow boundaries (Fig.5F). Further exploiting our cross-linker based models, we used analyses based on the radius of gyration and Mean First Encounter Time (MFET) to determine how TAD organization is influenced by boundary organization. Both analyses identified a considerable impact on the interface between TADs, which can extend far beyond the boundary region itself. Unexpectedly, we found that the addition of clustered CBSs at the boundary, represented by a gap in the model, strongly reduces both the number of monomers at the interface (Fig.6B and Fig.8F; particularly at more physiologically relevant *N_c_* numbers) and the interaction probability between loci at further distance from the boundary (Fig.7). This later aspect was mostly insensitive to a range of physiologically-relevant *N_c_* numbers, indicating that the formation of these inter-TAD contacts is a spurious process that does not require nearby blocking of loop extrusion. Reanalysis of Hi-C data confirms this observation, showing that extended TAD boundaries coincide with reduced inter-TAD contact between the neighboring TADs (Fig.8F).

### 2.1 Extended TAD boundaries, loop extrusion blocking and inter-TAD encounters

Cohesin and CTCF are essential for TAD formation through their intra-TAD loop extruding activity and blocking of this process at TAD boundaries, respectively (*e.g.* [53, 62, 63]). In this study, focusing on the width of the *IS* valleys at TAD boundaries, we report that the patterns of loop extrusion blocking are diverse and TAD boundary specific. Through the calibration of our cross-linker based polymer models, aimed at reproducing the *IS* pattern at each category of boundaries, we confirm the importance of different components of TAD boundary structure. Whereas blocking of loop extrusion is essential for all categories of boundaries—as may be expected from the strong enrichment of CBSs at TAD boundaries—we particularly identify the heterogeneity aspect as the strongest determinant to explain differences between boundary categories (Fig.3E-G). As such, the *IS* pattern around the boundary itself is mostly dependent on the size of the domain where loop extrusion blocking occurred with varying efficiency. We envision that narrow boundaries are characterized by a strong loop extrusion blocking activity within a small genomic interval, whereas at extended boundaries multiple instances of weaker blocking occur (*i.e.* with an increased chance for the loop extrusion machinery to read through the blocking site). Such a model is confirmed by our intersection of the TAD boundaries with other chromatin features, which reveal a highly punctuated and elevated enrichment at narrow boundaries and a more gradual and less pronounced enrichment within and around extended boundaries (Fig.4, Fig.5 and Fig.S4). The latter was particularly true for the Rad21 component of the Cohesin complex, which suggests not only a more spread out blocking of loop extrusion, but a generally reduced efficiency as well. Variable configurations of clustered CBSs, or other chromatin features with blocking capacity, thus will influence the insulating capacity and permeability of a TAD boundary, which may further modulate genome-associated functions [38, 61].

As an alternative explanation, we considered the possibility that the width of insulation could be determined by the chromatin environment, for instance due to the overall activity state of the chromatin (*e.g.* overlapping heterochromatin) or the distance from the neighboring boundaries. We therefore assessed TAD boundary overlap with Hi-C A/B compartments (A compartment: more transcriptionally active and euchromatin and B compartment: more transcriptionally inactive and heterochromatin), which form mutually exclusive and generally multi-Megabase domains in the Hi-C matrix [1]. Whereas we find a small but significant enrichment of extended TAD boundaries within the B compartment, for all categories the large majority of boundaries is present in the active A compartment (Fig.S8A). Similarly, the three categories of boundaries localize at comparable distances from their neighboring boundaries, with a slightly increased median distance for extended boundaries as well (490 kb for extended boundaries versus 400 kb for narrow and 420 kb for intermediate boundaries; Fig.S8B). TAD boundary width therefore is strongly influenced by the chromatin surroundings or the distance from the neighboring boundaries. Interestingly, we did notice an enrichment of narrow boundaries at the transition between A and B compartments, suggesting a potential function in separating euchromatin from heterochromatin (Fig.S8A). Due to the small number of boundaries in the narrow category, their total number at these transitions remains small relative to the other categories (n = 33 for narrow boundaries versus n = 61 and 63 for intermediate and extended boundaries, respectively).

Unexpectedly, our biopysical simulations reveal that the blocking of loop extrusion at (narrow) boundaries promotes inter-TAD contacts. This effect is observed both for loci close to the boundary (using our radius of gyration analysis; Fig.6B) and for loci at further distance (MFET analysis for loci at the equivalent of 250 kb from the boundary; Fig.7). In contrast, the effect was reverted when a wider array of blocking sites was included (*i.e.* by adding a gap between the TADs in our models). Although these inter-TAD encounters decrease when the density of cross-linkers increases (*i.e.* representing the density of extruded loops), the difference between boundaries without and with a gap remains relatively stable.

Despite the importance of loop extrusion blocking for the insulation at TAD boundaries, we deduce that these inter-TAD contacts at further distance from the the boundaries are of a more spurious nature and an indirect outcome from the process of loop extrusion. Blocking of the Cohesin complex by a CBS on one side switches the outcome of the loop extrusion process from bi-directional to uni-directional [24, 34]. Consequently, one-sided blocking of loop extrusion will cause a ’reeling in’ of the chromatin within the neighboring TAD towards the boundary (Fig.8H, left versus middle). In turn, loci within this TAD are brought into proximity of the neighboring TAD, thereby promoting the formation of spurious contacts with loci in the other TAD. The presence of multiple CBSs within extended boundaries, recently shown to improve the insulation between neighboring TADs through sequential loop extrusion blocking [38], may counter this effect by adding additional distance between the TADs at further distance from the boundary (Fig.8H, right). The analysis of Hi-C data confirmed that long-range inter-TAD contacts (≥ 200 kb) were increased around narrow TAD boundaries.

### 2.2 TAD boundaries versus TAD boundary interface

Boundaries are ubiquitous entities in sub-cellular biology, where they allow a regional delimitation to avoid protein or ion mixing (*e.g.* [64]). However, the term TAD boundaries is commonly used to indicate sites or zones in the linear genome where loop extrusion is blocked [2, 38, 53]. Our study aimed to determine how different types of boundaries within the (linear) genome affect the delimitation between neighboring TADs. To stay with common nomenclature, we use the term TAD boundary to refer to genomic regions dedicated to loop extrusion blocking. Instead, when discussing the insulated nature of TADs and their degree of intermingling, we refer to the (boundary) interface between TADs.

While the boundaries between cells and between the cytoplasm and the nucleus are membrane-based, the combined insulating effect of active intra-TAD loop extrusion and blocking at dedicated boundaries is much less stringent. Our models nonetheless identify variables that affect the interface between neighboring TADs, with unexpected distal influence of (linear) boundary composition. Using the radius of gyration and passage time analysis (MFET) as measures, we confirm a consistent influence of cross-linker number *N_c_*. In all models, an increased number of cross-linkers restricted the number of monomers in the region at the interface between the two TADS (Fig.6B). Similarly, the MFET analysis showed that more cross-linkers reduced the number of monomers around the linear boundaries whose behavior was affected (Fig.8F). Finally, we reported that the incorporation of extended boundaries, through the addition of a non cross-linked polymer, restricts the number of monomers at the interface and, unexpectedly, stabilizes the number of monomers with anomalous behavior. The latter is particularly prominent at lower *N_c_* numbers, which more closely mimic the numbers we used to replicate insulation at the different categories of TAD boundaries (Fig.3 and Fig.8F).

### 2.3 TADs boundaries as modulators of inter-TAD EP-looping

The perturbation of CTCF binding at CBSs can permit the formation of new EP-loops, thereby leading to ectopic activation in the context of embryogenesis and cancer [6, 7]. Our classification of TAD boundaries and MFET analysis indicate a dual impact on inter-TAD contacts: blocking of the active process of Cohesin-mediated loop extrusion at CBSs and the reduction of spurious TAD intermingling by extending the region where loop extrusion is blocked. Whereas loop extrusion promotes EP-loop formation at distances of ≥ 100 kb, it is not essential to permit the creation of stable and productive interactions between high-affinity enhancer-promoter pairs [65, 66, 67]. Spurious intermingling of neighboring TAD may thus be a non-negligible source of EP-loop formation, particularly in cases where developmental genes and enhancers on either side of a TAD boundary share mutual affinities (*e.g.* [54, 68]). The regulatory impact of clustered CBSs at TAD boundaries thus appears to extend beyond the blocking of loop extrusion alone, incorporating a possibility for fine-tuning of gene activity through the formation of more passive EP-contacts. Of note, all previously studied boundaries where arrays of CBSs contribute to the separation between neighboring TADs are part of the intermediate category of boundaries, suggesting these boundaries may be particularly optimized for the modulation of EP-loop formation of nearby developmentally regulated genes (Table S1 versus [38, 35, 36, 37, 68]). In contrast, our analysis of transcriptional deregulation upon CTCF depletion reveals an enrichment for genes in the direct vicinity of narrow boundaries, suggesting a more deterministic connection (Fig.4E). Expanding on the observed impact of local loop density, we envision that the combination of TAD boundary width and a modulation of local loop density will directly influence cross-boundary contacts both near the TAD boundary and at further distance. This may for instance be mediated through preferential loading of the Cohesin complex at regulatory elements or near CBSs [10, 38, 69, 70]. This mix of mechanisms can be particularly important for inter-TAD regulation of genes located close to a TAD boundary [71].

### 2.4 Are there limits to the width of TAD boundaries?

Our ranking of TAD boundaries revealed a wide range of widths, which raises the question if there is a limit to their seize. Based on the fitting of our RCL polymer models, we find that the range of values for most parameters is quite limited (*N_c_*, *N_gap_*, with the obligatory presence of fixed cross-linkers at theboundary). The most variable parameter appears to be the moving boundary position, which represents the heterogeneous nature of loop extrusion blocking. Our gyration radius and MFET analyses indicate that this parameter has little impact on TAD intermingling, either close to the boundary (Fig.6B) or at further distance (Fig.7B,C). More globally, the parameter does not noticeably influence the spreading of anomalous behaviour away from the boundary (Fig.8F). In contrast, the MFET analysis confirms the impact on inter-TAD encounters by the extended nature of the boundary and the blocking of loop extrusion. Yet, these parameters emerge as mostly invariant in the models that replicate the three boundary categories (Fig.3). Our strategy to infer the width of individual boundaries indicates that narrow and in intermediate boundaries are limited to ≈ 100 kb (Fig.5B). A potential explanation for this may be found in the mechanism for loop extrusion itself: wider boundaries may become susceptible to loading of Cohesin within and thus the formation of intra-boundary loops. Indeed, for both intermediate and extended boundaries we find a (moderate) enrichment of CBSs with a convergent orientation within the genomic interval covered by the boundary (Fig.5F), suggesting they may form chromatin structures that resemble TADs (see Fig.2A, right for an example). Since the process of loop extrusion and EP-loop formation are closely linked ([54, 66, 67, 69] and this study), we envision that these boundaries may have unrecognized functions in gene regulation. Future studies that incorporate the linear organization of blocking sites at individual boundaries, together with the analysis of 3D organization and inter-TAD insulation, may help to identify such cases of ’super-extended’ boundaries with gene regulatory functions.

## 3 Materials and Methods

### 3.1 Reanalysis of published Hi-C

Hi-C data from mouse ESCs, NPCs and CNs were reused from a previous study [44], using a previously described strategy [38]. Raw paired-end reads were obtained from https://www.ncbi.nlm.nih.gov/ geo/query/acc.cgi?acc=GSE96107, followed by mapping to the ENSEMBL Mouse genome assembly GRCm38.p6 (mm10) and further processing using the HiC-Pro pipeline (v2.9.0) [72] to obtain an ICE (Iterative Correction and Eigenvector decomposition) normalized interaction matrix at 10kb resolution. A/B compartments were calculated using the cooltools API (v0.5.2) [73], with manual orienting of the eigenvector sign per chromosome based on gene content.

#### 3.1.1 Insulation score (*IS*) and TAD boundaries

Insulation score (*IS*) and TAD boundaries for mESCs were computed using TADtool (v0.76) [74] with a previously used window size of 500kb and a cutoff of 21.75. To avoid bias due to difference in genome coverage, all downstream analysis was limited to the autosomes, without including the X and Y chromosomes. The list of TAD boundaries used in this study is provided in Table S1. The first and second derivative of the *IS* (*IS’* and *IS”*) were computed from the raw *IS* or the first derivative of the *IS* by subtracting each 10kb bin value from its downstream bin.

For an improved comparison between the *IS* and *IS”* between mESCs, mNPCs and mCNs (Fig. S5A), Hi-C matrices were normalized using the Cooler tool (v0.5.2) [73]. The *IS* was subsequently determined from the resulting matrices in the cool-format using the FAN-C tool (v0.9.1) [75]. Although TADtool and FAN-C globally returned similar patterns for the *IS*, we noticed increased variability in FAN-C signal that complicated the reliable inference of boundary width (see below). In spite of the need for normalization between samples, we nonetheless preferred this tool for our comparative analysis. For all analysis, the TAD boundaries as initially identified from the TADtool analysis were used.

Heatmaps and pileups for the relative *IS* and its derivatives over a chosen flank distance were computed and plotted using the computeMatrix and plotHeatmap commands from deepTools2 (v3.5.1) [76]. In these graphs, the boundary bin is at the center, with an equal number of bins upstream and downstream. For all boundaries, the center bin for the *IS* was first normalized to 0 by running the computeMatrix and plotHeatmap commands to extract the raw *IS* matrix in a usable format. Next, a second call to the plotHeatmap command was performed to obtain final plots.

TAD insulation relative to distance and boundary categories were determined using a previously published approach [3]. Read pairs spanning 10 kb - 1 Mb were categorized in four categories based on the absence of a boundary or the presence of one of the three categories of TAD boundaries in-between. Read pairs separated by more than one boundary were removed. Median interaction signal for all read pairs in each category was calculated based on the distance.

#### 3.1.2 Classification of TAD boundaries using a K-means algorithm

K-means clustering into three clusters has been performed on the second derivative of the *IS* using the `--kmeans 3` option of the plotHeatmap command of the deepTools2 suite (v3.5.1) [76]. The same clusters and order of boundaries were used in all plots. The list of boundaries included in the three categories is provided in (Table S1). Average Hi-C matrices for the different clusters were computed using the cooltools API (v0.5.2) [73] (cooltools.pileup with a 250kb flank). Half of the resulting symmetric matrix is displayed. A/B compartments were computed using the cooltools CLI (v0.5.2) (cooltools eigs-cis command) and compared using R (v4.2.2) with Bioconductor (v3.17) [77] and package BRGenomics (v1.12.0) [78].

#### 3.1.3 Reanalysis of published ChIP-seq, ATAC-seq, PRO-seq and RNA-seq data

For the average enrichment of chromatin features arpund TAD boundary categories, CTCF and Rad21 ChIP-seq data from mESCs were reused from [48, 38] using a previously described strategy [38]. Briefly, unprocessed data were obtained from https://www.ebi.ac.uk/ena/browser/view/PRJEB44135 and https://www.ncbi.nlm.nih.gov/geo/query/acc.cgi?acc=GSE33346, followed by mapping to the ENSEMBL Mouse genome assembly GRCm38.p6 (mm10) using BWA (v0.7.15) [79] with default parameters. ATAC-seq data from mESCs were reused from a previous study [51]. Unprocessed data were obtained from https://www.ncbi.nlm.nih.gov/geo/query/acc.cgi?acc=GSE138923, followed by mapping using bowtie2 (v2.5.1) [80] and processing using SAMtools (v1.17) [81] and deepTools2 (v3.5.1) [76]. PRO-seq data from mESCs were reused from a previous study [52]. Unprocessed data were obtained from https://www.ncbi.nlm.nih.gov/geo/query/acc.cgi?acc=GSE130691, followed by mapping to the ENSEMBL Mouse genome assembly GRCm38.p6 (mm10) and further analysis using the proseq2.0 pipeline [82] obtained from https://github.com/Danko-Lab/proseq2.0/. For all datasets, the ENCODE blacklist for mouse genome assembly GRCm38.p6 (mm10) was used (available from https://github.com/Boyle-Lab/Blacklist/blob/master/lists/mm10-blacklist.v2.bed.gz).

For the preparation of histograms, matrices were computed consisting of mean values in 10kb bins around each TAD boundary using deepTools2 computeMatrix (v3.5.1) [76]. A maximum threshold of 1000 (`--maxThreshold 1000` in the computeMatrix command) was applied to all values, in order to remove outliers. Genome-wide average signal for ChIP-seq and PRO-seq datasets was determined using the `meanI` command from WiggleTools (v1.2) [83] on the bigWig files. For the ATAC-seq dataset, mean values in 10kb bins were computed using the bedtools suite [84], followed by filtering of bins that overlap blacklisted regions. The resulting bedGraph file was converted to the bigWig format using the bedgraphtobigwig tool (available from http://hgdownload.soe.ucsc.edu/admin/exe/). Genome-wide average signal was subsequently determined as described for ChIP-seq and PRO-seq data.

Lists of significant ChIP-seq CTCF binding peaks in mESCs, mNPCs and mCNs (used in Fig. S5A) were reused from [44] (obtained from https://www.ncbi.nlm.nih.gov/geo/query/acc.cgi?acc=GSE96107). For the enrichment of deregulated TSS around boundaries, a list of deregulated genes in CTCF-AID mESCs was obtained from Table S3 in [53]. Filtering of gene symbols against the BioMart GRCm38.p6 (mm10) Ensembl 102 gene list (http://nov2020.archive.ensembl.org/Mus_musculus/Info/Index) returned 21988 genes on the autosomes, where we assigned the start coordinate of each gene as the TSS. Among the 21988 TSS, 662 were assigned as upregulated after 2 days of Auxin treatment and 579 were assigned as downregulated [53]. TSS were assigned to 10 kb bins in the genome, followed by filtering against TAD boundaries in the different categories.

#### 3.1.4 Inference of width for individual boundaries from the IS”

To obtain a measure for the width of individual boundaries, we developed a strategy using on thresholding of the normalized second derivative of the insulation score (*IS”*) around identified TAD boundaries. The normalizerd deepTools2 [76] matrix files for the *IS”* in 250 kb windows up- and downstream of the boundaries (51 bins), used for the computing heatmaps and pileups, were taken as input. First, we determined the average *IS”* value for all fields in the matrix (all 51 bins for the 3654 identified boundaries), which served as our cut-off value. For each individual boundary, taking the central bin (which was normalized to 0) as reference point, we next determined the first bin up- and downstream with a value below the average *IS”* value. Boundary width for each individual boundary is subsequently calculated by taking the boundary bin and the bins on both sides up to (but not including) bins with values below the average. Subsequently, two additional steps of filtering were implemented. First, if the normalized *IS”* value on both sides of the central bin had average values below the cut-off (returning a width of 1 bin), we included the bin directly on the left, as we noticed the first derivative of the *IS* (*IS’*) passed the 0 value in this bin in a considerable fraction of boundaries. Second, if the *IS”* value in the 25 bins up- or downstream of the boundary did not contain any values below the cut-off, we assigned the ”undefined” status to the boundary. The inferred range for individual boundaries is provided in Table S1. To determine the distribution of CTCF peaks, peak signal and motif orientation, a detailed list of CTCF peaks in mESCs was obtained from Supplementary Data file 1 in [38]. Combined peaks were assigned to 10 kb bins in the genome, with only the orientation of the most significant motif retained for each peak, and for bins with multiple peaks, their signal combined and the orientation of all peaks retained. For all boundaries where we could assign a defined width, CTCF features were extracted for the two bins at the extremities of the boundary and the 5 bins up- and downstream of those bins. For boundaries that covered 3 bins or more, CTCF features in the internal bins were proportionally divided over 2 bins (narrow boundaries) or 8 boundaries (intermediate or extended bins). For all bins, signal for all boundaries in each category was combined after normalization for the total number of boundaries in the category.

### 3.2 Modeling and accounting for TAD boundary components

To account for the different types of boundary between two subsequent TADs, we use a block polymer of *N_mon_* monomers composed by two Random Cross-Linked polymers (RCL) [25, 28]. Each of these is composed by 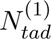 and 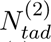 monomers respectively. Monomers belonging to each of the two TADs have indexes 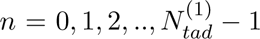 and 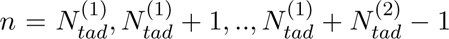 respectively, we consider only random connector distributions such that only monomers belonging to the same TAD are connected. To investigate how the non-uniform distribution of random connectors affects the boundary between the two RCL polymers, we added a dictionary of possible connectivity:

1. **Uniform distribution of random connectors** within the same block (TAD) of the polymer where 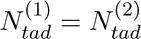;
2. **A gap of** *g* **monomers without any random connectors**, separating two identical RCL blocks, such that 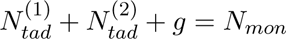 and 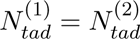;
3. **A random boundary position shift**, where 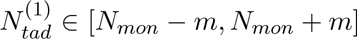 is a random integer number with *m >* 0 and 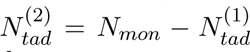 and random connectors can join only monomers belonging to the same block;
4. **Enriched connectivity at the boundary:** this is realized by enforcing the presence of two random connectors with one side lying on the last monomer before the boundary for each of the two TADs.

The dynamics of the polymer in the solvent is described by the over-dumped limit Langevin’s equation

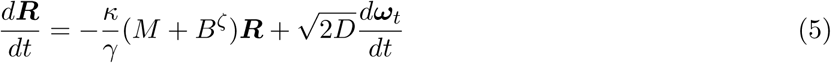

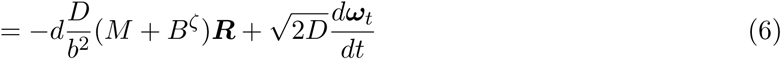

where 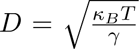 is the diffusion coefficient, *T* the temperature, *γ* the friction and *ω_t_* is the standard d-dimensional Gaussian noise with average 0 and standard deviation 1. A random connector joins two monomers *n*, *m* randomly-chosen (where |*n* − *m*| *>* 1). Given a polymer realization, defined by a set *ζ* of *N_c_* random connectors, the interaction among monomers is determined by the interaction matrices *M*, sum of a Rouse matrix [14]

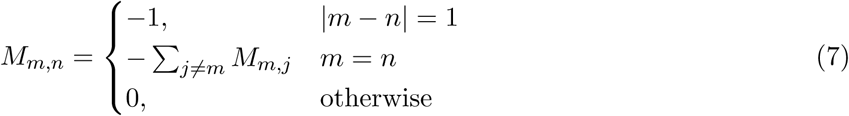

and the *B^ζ^*matrix is defined from the ensemble *ζ* of connectivity interactions:

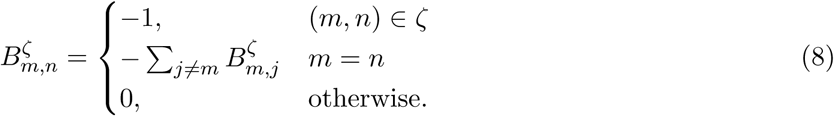

In general, we considered polymer composed by *N_mon_* = 200 monomers, with *N_tad_* = 100 monomer for each TAD. The number of monomers in a gap varies in the range of *N_gap_* ∈ [0, 10] and *N_var_* ∈ [0, 10]. The polymer parameters are given by *b* = 0.18*µm*, *ɛ* = 0.06*µm*, with a diffusion coefficient *D* = 8 · 10*^−^*^3^*µm*^2^*/s* [25, 38].

Numerical simulations are performed by integrating the Langevin’s equation 5 with a time step Δ*t* = 10*^−^*^2^*s* first for 10^6^ integration steps to reach equilibrium, then for 10^6^ to collect the configurations to compute the needed statistics reported in this manuscript. Finally we continue the simulations up to maximum of 10^8^ integration steps to find the mean first encounter time. Ensemble averages are performed over 10^3^ polymer realizations.

## Acknowledgments

This work benefited from collaborative funding from PlanCancer (19CS145-00) and the Agence Nationale pour la Recherche (ANR-24-CE12-5087) to Daan Noordermeer and David Holcman. Work in the Holcman lab is further supported by funding from the Agence Nationale pour la Recherche (AnalysisSpectralEEG, AstroXcite, Memolife) and the European Research Council (ERC) under the European Union’s Horizon 2020 research and innovation programme (No 882673). Work in the Noordermeer lab is further supported by funding from the Agence Nationale pour la Recherche (ANR-21-CE12-0034). Andrea Papale was supported by a postdoctoral fellowship from the Fondation pour la Recherche Médicale (SPF201909009284). Julie Segueni was supported by a PhD grant from La Ligue contre le Cancer (Allocations Doctorales 4èeme Année de Thèese).

## Data and codes availability

Codes and data are available at https://zenodo.org/records/14232689

