## Supplementary material for "Insulation between adjacent TADs is controlled by the width of their boundaries through distinct mechanisms": SI

[+,@] equal contributions.

December 25, 2024

### Supplementary figures

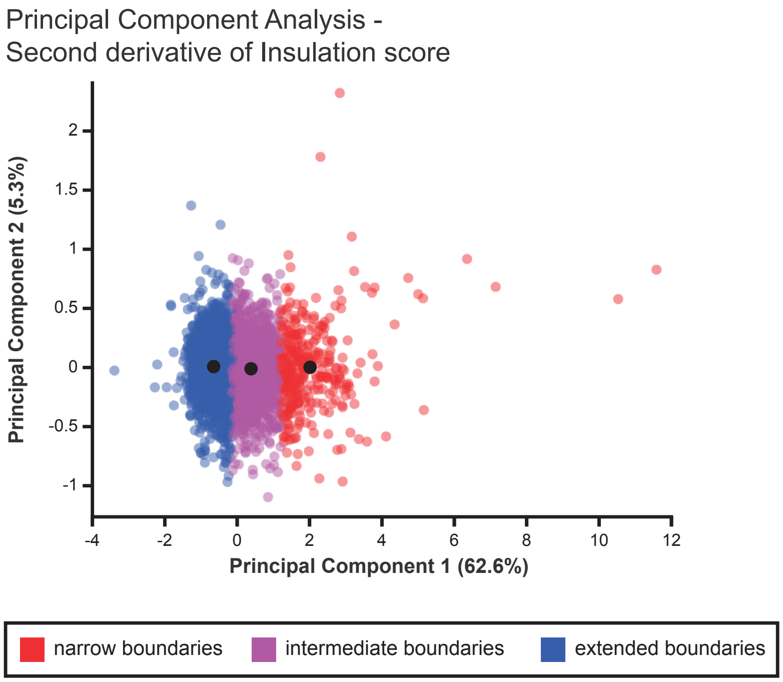


Figure S1: **Principal component analysis of TAD boundaries with different widths.** Visualization of the first and second principal component of the second derivative of the insulation score ($IS''$) for the three boundary clusters as described in Fig. 2, which together make up nearly 68% of the variance. Although the contribution of the additional principal components is taken along in the $k$-means clustering approach as well, the three clusters already show a nearly complete separation on the x-axis.

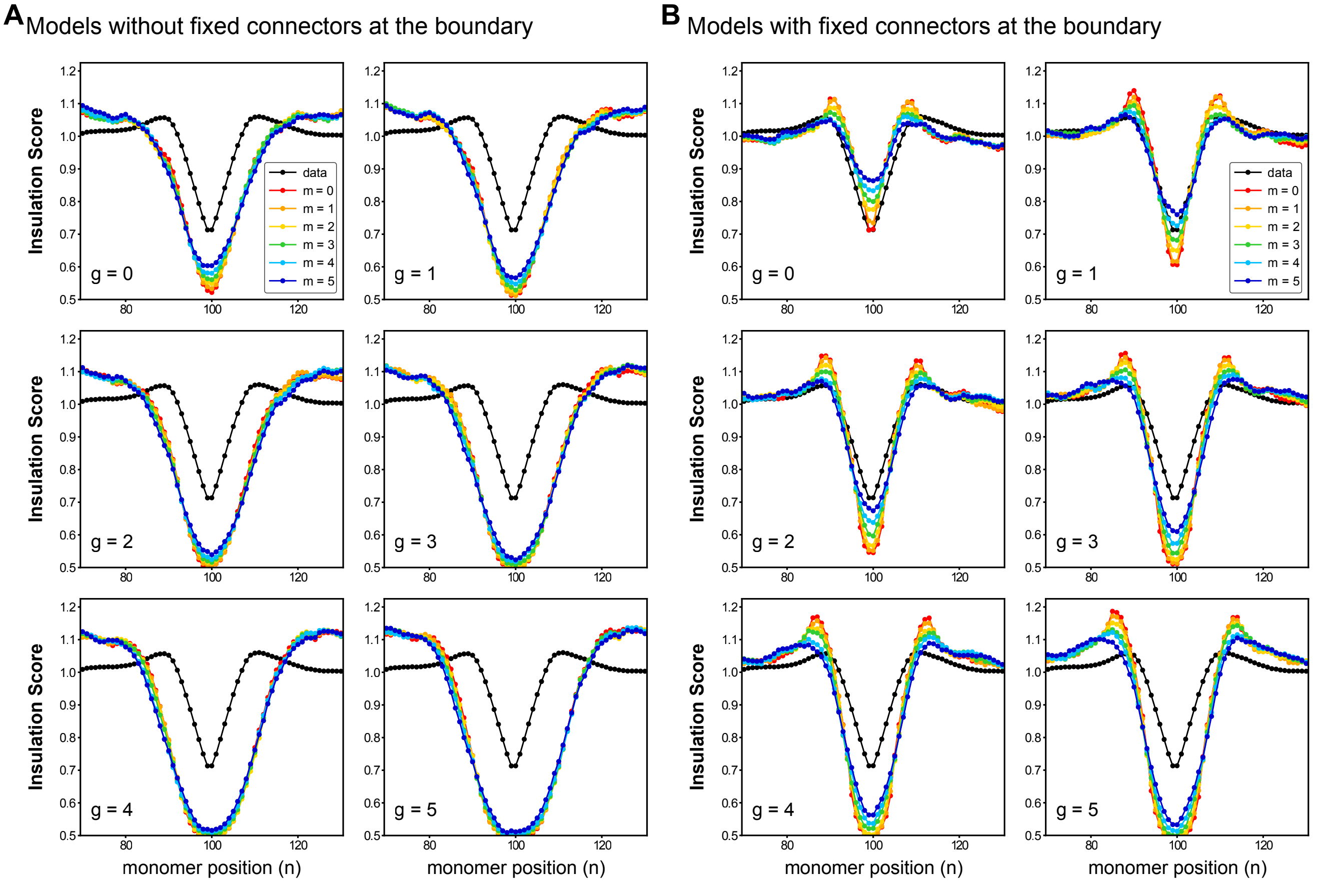


Figure S2: **Insulation scores for various parameter values.**
Calculation of the two-dimensional insulation score ($IS$) vs monomer location, using a wide-range of parameters. **(A)** $IS$ for models without a fixed connector at the boundary. **(B)** $IS$ for models with a fixed connector at the boundary. Different panels represent the $IS$ with an $N_{gap}$ of different monomers at the boundary ($N_{gap}$ = 0-5). Colored lines represent different values for the moving boundary position (variable $m$ = 0-5). The black line indicares the average genome-wide $IS$ at TAD boundaries, as obtained from Hi-C experiments.

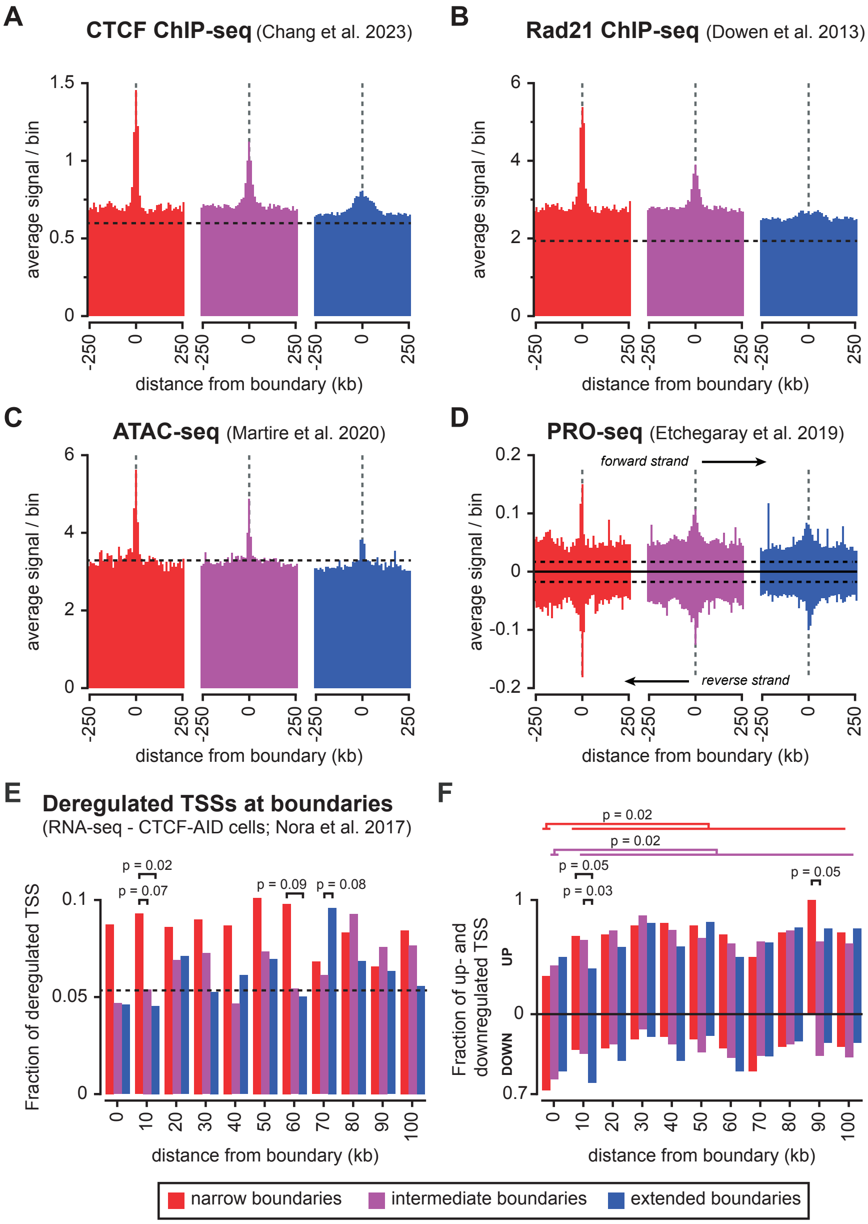


Figure S3: **Enrichment of chromatin and gene regulatory features in extended regions around the three categories of TAD boundaries.**
**(A)** Average distribution of CTCF ChIP-seq signal. **(B)** Average distribution of Rad21 ChIP-seq signal. **(C)** Average distribution of accessible chromatin signal (ATAC-seq). **(D)** Average distribution of transcription signal (PRO-seq), with signal separated on the plus and minus strands. **(E)** Fraction of deregulated Transcriptional Start Sites (TSSs) among all TSSs within bins upon CTCF depletion (RNA-seq in 2-days Auxin-treated CTCF-AID cells). **(F)** Fraction of up- and downregulated TSSs among deregulated TSSs within bins upon CTCF depletion. The origin of the datasets, all from mouse ESCs, is indicated above each panel. Signal is binned to 10 kb, with dashed lines representing the average genome-wide enrichment. Signal in panels **(A-D)** is shown in a window of 250 kb up- and downstream of TAD boundaries. Signal in panels **(E,F)** is merged on both sides of the TAD boundaries and shown in a window up to 100 kb away from the boundaries. Significance of difference between categories for the same bin [panel **(E,F)**] and for the 0 kb bin versus the combined 10-100 kb bins within the same category [panel **(F)**] were determined using a G-test of independence.

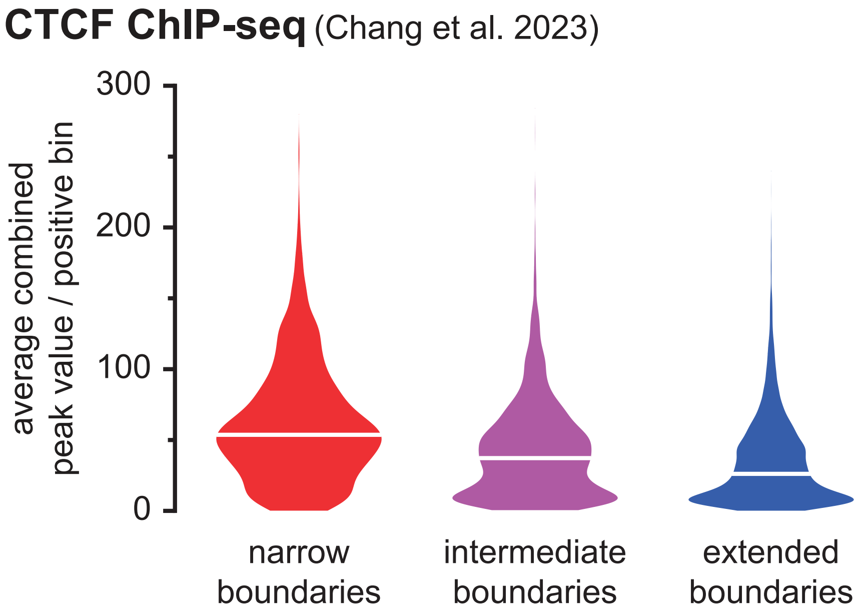


Figure S4: **Distribution of CTCF peak signal within inferred boundaries.**
Distribution of CTCF ChIP-seq signal (combined signal of all peaks in 10 kb bins) within the inferred boundaries in mouse ESCs for the different boundary categories.

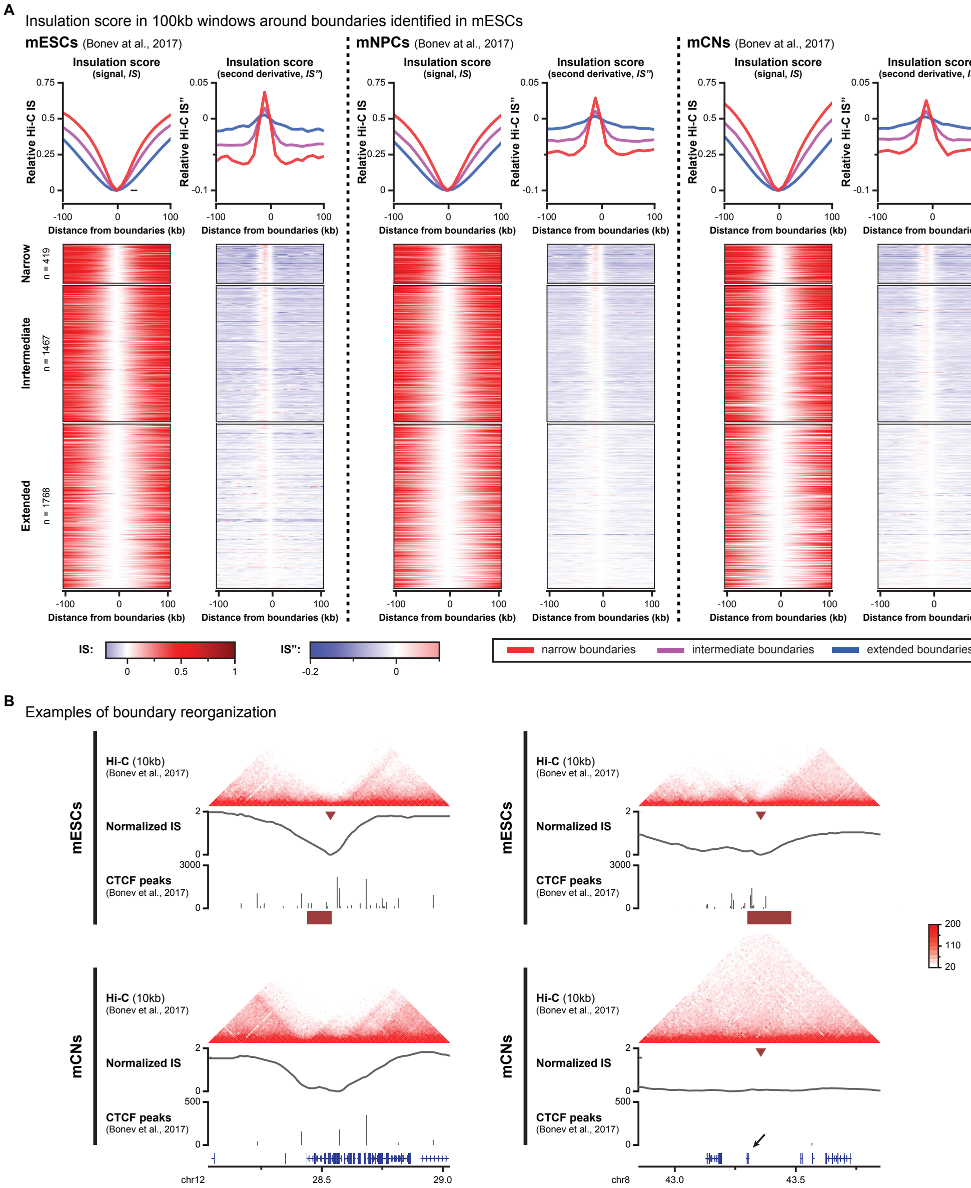


Figure S5: **Developmental dynamics of boundary width.**
**(A)** Pile-ups and heatmaps of the normalized insulation score *IS* and its normalized second derivative (*IS"*) in mESCs, Neural Progenitor Cells (mNPCs) and Cortical Neurons (mCNs) around the TAD boundaries previously identified in mESCs. Average signal for the three categories in a window of 100 kb up- and downstream of the boundaries is indicated on top and heatmaps for individual boundaries are depicted below. **(B)** Examples of TAD boundaries with changes in insulation width between mESCs and mCNs, associated with reorganization of CTCF binding. Hi-C data is depicted above. Purple arrowheads indicate the position of the boundary in mESCs. Below, tracks for the Hi-C insulation score, CTCF peaks and computationally inferred width of the boundaries (purple boxes are shown. The arrow in the gene track for the right example indicates the *Zfp42* gene, a pluripotency marker (also known as *REX-1* in the human genome), whose activity is restricted to embryonic stem cells.

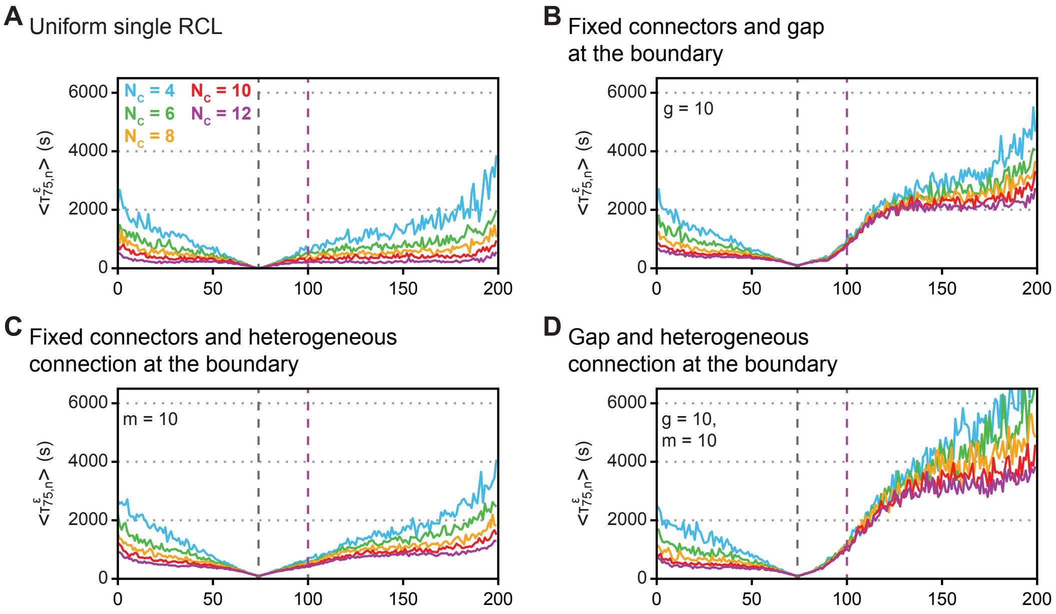


Figure S6: **MFET for additional connectivities**

**(A)** MFET for a single RCL (*i.e.* a single polymer without internal separation) with different numbers of random connectors $N_c = 4, .., 12$ obtained averaging over $10^3$ polymer realizations. The black dotted line indicates the position of monomer 75. The purple dotted line indicates monomer 100, which is the the boundary between TAD 1 (left) and TAD 2 (right) in the other connectivities. **(B)** MFET for a connectivity that combines a fixed connector and a gap at the boundary. **(C)** MFET for a connectivity that combines a fixed connector and a a heterogeneous connection at the boundary. **(D)** MFET for a connectivity that combines a gap and a a heterogeneous connection at the boundary.

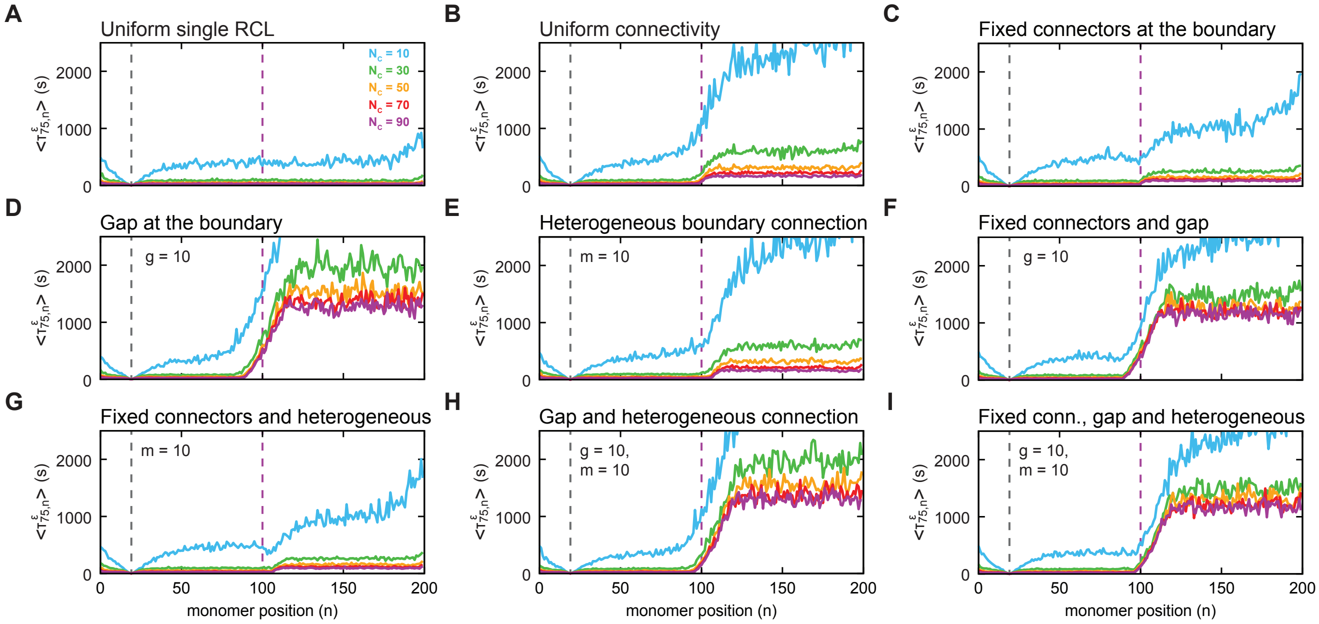


Figure S7: **MFET over larger distance and with increased numbers of cross-linkers.** **(A-I)** MFET for the different connectivities as presented in Fig. 7 and Fig. S6, but with increased numbers of random connectors ($N_c = 10, .., 90$) and using monomer 25 as anchor. MFET was obtained averaging over $10^3$ polymer realizations. The purple dotted line indicates monomer 100, which is the the boundary between TAD 1 (left) and TAD 2 (right) in the other connectivities.

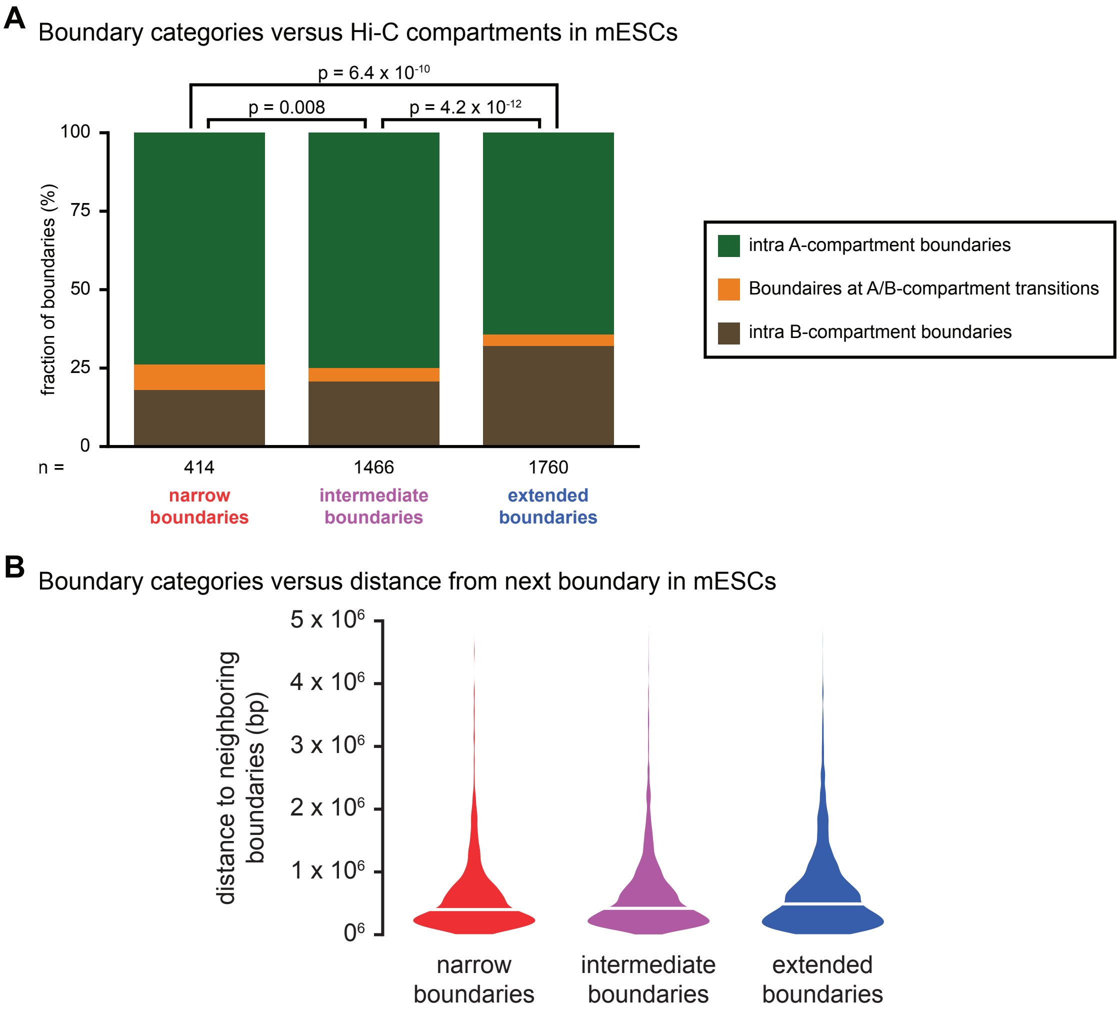


Figure S8: **Overlap of TAD boundaries with A/B compartments and distance from neighboring boundaries.**
**(A)** The location of TAD boundaries was scored relative to the position of Hi-C A/B compartments. The orange category represents TAD boundaries that overlap a transition between the A and B compartment. Significance of difference between the three categories was determined using a G-test of independence. Narrow boundaries are relatively enriched at the transition between A/B compartments, although the small size of this category means the number remains limited. Extended boundaries are relatively more enriched within the B-compartment. **(B)** Distribution of distances from neighboring boundaries for the different boundary categories. For all boundaries, both the distance to the nearest up- and downstream boundary was included.
